# Chimpanzee pant-hoots encode information about individual but not group differences

**DOI:** 10.1101/2021.03.09.434515

**Authors:** Nisarg P. Desai, Pawel Fedurek, Katie E. Slocombe, Michael L. Wilson

## Abstract

Vocal learning, the ability to voluntarily modify the acoustic structure of vocalizations based on social cues, is a fundamental feature of speech in humans (*Homo sapiens*). While vocal learning is common in taxa such as songbirds and whales, the vocal learning capacities of nonhuman primates appear more limited. Intriguingly, evidence for vocal learning has been reported in chimpanzees (*Pan troglodytes*), for example in the form of regional variation (‘dialects’) in the ‘pant-hoot’ calls. This suggests that some capacity for vocal learning may be an ancient feature of the *Pan-Homo* clade. Nonetheless, reported differences have been subtle, with inter-community variation representing only a small portion of the total acoustic variation. To gain further insights into the extent of regional variation in chimpanzee vocalizations, we performed an analysis of pant-hoots from chimpanzees in the neighboring Kasekela and Mitumba communities at Gombe National Park, Tanzania, and the geographically distant Kanyawara community at Kibale National Park, Uganda. We observed group differences only among the geographically isolated communities and did not find any differences between the neighboring communities at Gombe. Furthermore, we found differences among individuals in all communities. Hence, the variation in chimpanzee pant-hoots reflected individual differences, rather than group differences. The limited evidences for vocal learning in *Pan* suggest that extensive vocal learning emerged in the human lineage after the divergence from *Pan*.

## Introduction

Vocal learning underlies the human capacity for speech. The desire to understand the evolution of this capacity motivates much of the research into vocal learning in other animals (Fitch, 2010). Over time, the definition of vocal learning has evolved as researchers have identified several nuances in vocal learning ability across animals. Janik & Slater (2000) defined vocal production learning broadly, as “signals modified in form as a result of experience with those of other individuals.” Other researchers have focused on more specific aspects, such as the ability to voluntarily modify and learn new vocalizations through imitation (Fitch, 2010). Regardless of the particular definition used, it is clear that vocal learning has evolved independently multiple times in animals. For example, songbirds (Passeriformes) and humpback whales (*Megaptera novaeangliae*) learn elaborate songs (Cunningham & Baker, 1983; Garland et al., 2011); parrots (Psittaciformes) can mimic human speech, and distinguish group members from drifters based on learned vocalizations (Bartlett & Slater, 1999; Hile & Striedter, 2000). In comparison to birds and whales, the vocal learning capacities of nonhuman primates appear much more limited (Fischer & Hammerschmidt, 2020). Researchers generally agree that non-human primates are unable to actively learn new vocalizations (Tyack, 2020), although recent studies indicate that orangutans (*Pongo spp.*) can acquire and produce novel vocalizations (Lameira et al., 2015; Wich et al., 2009). Some non-human primates have been reported to subtly modify acoustic structure of vocalizations based on auditory feedback and imitation. Takahashi and colleagues (2015) found that common marmosets (*Callithrix jacchus*) learn vocalizations through parental feedback (Takahashi et al., 2015). Zurcher & Burkart (2015) observed that common marmosets exhibit dialects in their vocalizations. Sugiura (1998) reported that Japanese macaques (*Macaca fuscata*) match some of the acoustic features of recorded coo calls during a playback experiment. Fischer et al. (2020) found that the grunts of male Guinea baboons (*Papio papio*) that interacted more frequently with one another exhibited greater resemblance than the grunts of males that interacted less frequently.

Much of the literature on vocal learning in animals focuses on regional variation in vocal production: dialects. When such variation is learned, it may signal membership in the local population (as in songbirds (Cunningham & Baker, 1983)), or membership in a particular social group (as in wolves, orcas (Filatova et al., 2012; Zaccaroni et al., 2012)). Studies of social birds and mammals have found that learned signals of group membership can benefit individual signalers in various ways. Comparison with other species suggests two main ways in which chimpanzees might benefit from producing learned signals of group membership: (i) by eliciting affiliative interactions from group members and mates and/or (ii) by advertising group membership to rivals during agonistic interactions, such as during territory defense. For example, in birds, group-specific calls can (i) help maintain social bonds among group members or help choose mates, as in budgerigars (*Melopsittacus undulatus*) (Farabaugh et al., 1994; Hile & Striedter, 2000); (ii) facilitate territory defense by helping individuals distinguish flock members from drifters and prevent potential agonistic interactions during inter-flock encounters, as in black-capped chickadees, *Parus atricadpillus* (Nowicki, 1983). Researchers have inferred similar functions in social mammals. For example, several species of toothed whales (Odontocetes) appear to use vocal dialects to facilitate spatial group cohesion and maintain social relationships (Janik, 2014; Tyack & Sayigh, 1997). Spatial cohesion is necessary in group living species to help maintain social bonds, find mates, or to defend territories (Janik & Slater, 1998). Several canid species appear to use group-specific howls to facilitate spatial group cohesion, as well as to defend territory (Kershenbaum et al., 2016).

Dialects have figured prominently in efforts to find evidence of vocal learning in chimpanzees (*Pan troglodytes*) (Fedurek & Slocombe, 2011). Researchers interested in the origins of human language have been particularly interested in the vocal behavior of chimpanzees (*Pan troglodytes*), given that they are one of the two living species most closely related to humans (Fedurek & Slocombe, 2011). Several studies from the field (Arcadi, 1996; Crockford et al., 2004; Mitani et al., 1992) and captivity (Marshall et al., 1999) have found evidence for regional variation in chimpanzee pant-hoot calls, which has been proposed to result from vocal learning (Crockford et al., 2004; Marshall et al., 1999). Pant-hoots of males that spend more time together are more similar, and the acoustic features of their calls converge when chorusing together (Mitani & Brandt, 1994), suggesting a possible mechanism for the convergence of acoustic properties within groups (Fedurek, Schel, et al., 2013; Mitani & Gros-Louis, 1998). Call convergence has also been reported for chimpanzee rough-grunt calls (Watson et al., 2015). Similar to wolves (*Canis lupus*) and other canids, chimpanzees live in groups with fission-fusion dynamics, in which individuals travel in subgroups (known as ‘parties’) of varying size, and they communicate over long distances using vocalizations, often in noisy environments. Thus, vocal dialects potentially facilitate spatial group cohesion, and territorial defense during intergroup encounters.

Pant-hoots are structurally complex, with considerable variation within and among individuals in the acoustic structure (Fedurek, Schel, et al., 2013; Marler & Hobbett, 1975). Chimpanzee pant-hoots have a relatively consistent temporal patterning, which typically involves a sequence of four kinds of sound elements over a duration range of 2-20s. Each sequence of similar elements is called a phase and so the pant-hoots have four main phases (see Methods for details). The variation is not limited to frequency properties of elements such as fundamental frequency, peak frequency, etc. but also involves variation in the number and presence/absence of different elements and phases. Chimpanzees use pant-hoots in a variety of intra-community and inter-community contexts. In intra-community contexts, chimpanzees use pant-hoot calls to communicate with members of their own community over long distances (Goodall, 1986). Pant-hoots may function to communicate the caller’s location to allies and associates within their own community (Goodall, 1986; Mitani & Brandt, 1994). Further, pant-hoots play a role in facilitating social bonds as affiliative partners chorus more together (Fedurek, Machanda, et al., 2013) and play a role in regulating grouping dynamics by attracting allies and potential mates to the caller’s location (Fedurek et al., 2014; Mitani & Nishida, 1993; Wrangham, 1977). In inter-community contexts, interactions often involve hearing — and sometimes responding to — pant-hoots from callers that are hundreds of meters away, far out of view (Wilson et al., 2012). The long-distance nature of pant-hoots allows chimpanzees to use pant-hoots to advertise territory ownership (Wilson et al., 2007), and to signal numerical strength to members of neighboring communities during agonistic intergroup encounters (Herbinger et al., 2009; Wilson et al., 2001, 2012). Individual callers might thus benefit from encoding community-specific cues. Playback experiments have demonstrated that chimpanzees can distinguish stranger pant-hoots from those of familiar individuals (Herbinger et al., 2009) and that they are sensitive to numerical strength during intergroup encounters, being more likely to respond to simulated intruders when they are in parties with more males (Wilson et al., 2001). Hence, community-specific dialects could play a role in cooperative defense by signaling community membership. Insofar as genetics affect the acoustic structure of signals, socially learned signals of group membership might be useful in cases where not all group members are close kin.

Despite these reasons for thinking that vocal dialects would benefit chimpanzees, current evidence raises several questions about the extent to which chimpanzees have socially learned signals of group membership. In the first study of chimpanzee dialects, Mitani and colleagues reported differences between Gombe and Mahale pant-hoots and suggested that they may be an outcome of vocal learning (Mitani et al., 1992). However, the differences among the communities were subtle compared to differences observed in songbirds (Cunningham & Baker, 1983) or whales (Garland et al., 2011), with geographical differences in the composition identified for only one of the four main phases of the pant-hoot call, the build-up, and in frequency properties of another phase, the climax (Mitani et al., 1992). Mitani & Brandt (1994) later found that community membership accounted for only 11% of the variation in acoustic structure, with most of the remaining variation being due to variation within individuals. Mitani further reassessed his findings, pointing out that since Gombe and Mahale are far from one another (∼160 km) and likely genetically isolated, the acoustic differences may not necessarily represent vocal learning, but instead could represent genetic differences and/or body size (Mitani et al., 1999). Additionally, other environmental factors like habitat acoustics and/or sound environment might be more important in explaining the variation in such geographically distant communities.

In addition to assessing whether pant-hoots signal group membership, researchers have studied the acoustic structure of pant-hoots produced different contexts, such as traveling, feeding, group fusion, arrival at food sources (Clark & Wrangham, 1993, 1994; Fedurek et al., 2016; Goodall, 1986; Mitani & Nishida, 1993; Notman & Rendall, 2005; Uhlenbroek, 1996; Wrangham, 1977). Clark & Wrangham (1993) and Fedurek et al. (2016) observed an association of some properties of the letdown phase of the pant-hoots with the context in which the pant-hoot was produced. Notman & Rendall (2005) and Uhlenbroek (1996) observed an association of the tonal structure of the climax scream element of the pant-hoots with the context of the production. While this variation provides information about context to receivers, Notman & Rendall (2005) argue that these differences are unlikely to be an outcome of vocal learning and are more likely to reflect arousal states of chimpanzees when calling. In any case, the context of the call production is a potentially confounding factor that should be controlled for when testing for group differences. Finally, as Marler & Hobbett (1975) noted previously, pant-hoots are individually distinctive. Signaling individual identity, rather than group membership, might therefore be the primary function of these calls.

To test the extent to which the acoustic structure of pant-hoots specifically signals community membership and arises out of vocal learning via auditory feedback, three questions need to be answered: (i) Do the calls contain reliable acoustic cues of community membership that might allow chimpanzees to distinguish extra-community pant-hoots based on those cues alone, rather than through familiarity with the calls of particular individuals? (ii) Do chimpanzees from neighboring communities have more distinct pant-hoots than those from geographically distant communities? Greater differences among neighboring communities compared to geographically distant communities would indicate that chimpanzees are actively modifying the acoustic structure of pant-hoots to differentiate their calls from those of neighbors. (iii) Do genetically close kin have more similar calls? Crockford et al. (2004) addressed all three of these questions by comparing genotyped individuals in three neighboring communities and one more distant community in Taï National Park, Côte d’Ivoire. They found that neighboring communities differed from one another more than they differed from the distant community, supporting the view that chimpanzees learned to produce an acoustic structure distinct to their own community. This study thus supports the view that vocal learning accounts for the acoustic differences among communities. However, sample sizes in this study were small, with calls from only three individuals per group analyzed, raising the possibility that the findings are a statistical artifact resulting from small sample size. It is a common assumption that in situations where data are likely to be noisy (as is common in animal behavior studies), the observed effect sizes would be smaller than actual effect sizes. And based on this assumption, statistically significant results from noisy data are often inferred to be strong evidence of an underlying effect. However, this is a false assumption, especially when the sample sizes are small (Loken & Gelman, 2017). A small sample size with noisy data, such as that in Crockford et al. (2004), could artificially exaggerate effect sizes (*ibid*.). Hence, a larger sample of individuals is needed to be confident that differences in acoustic structure among communities are not the result of chance.

As a step towards reevaluating the role of vocal learning in chimpanzee calls, we recorded pant-hoot calls from two neighboring chimpanzee communities in Gombe National Park, Tanzania. To provide context for population-level differences, we compared calls recorded in Gombe with calls recorded from the geographically distant Kanyawara community of chimpanzees in Kibale National Park, Uganda. The objective of the study is to assess the extent to which variation in the acoustic structure of the pant-hoots can be explained by community membership. To that end, we test two hypotheses. Our first hypothesis is: the acoustic structure of pant-hoots contains socially learned elements that provide reliable cues of community membership. The second hypothesis is: the acoustic structure of pant-hoots contains cues of individual identity more than community identity. If there are community differences in the acoustic structure of the pant-hoots after adjusting for individual and context differences, it may indicate the possibility that these differences arise due to vocal learning. While these are not mutually exclusive hypotheses (i.e. one or both or neither could be supported), they provide a relevant framework to study our research questions.

## Methods

### Subjects and study site

We studied chimpanzees at two study sites: Gombe National Park, Tanzania and Kibale National Park, Uganda. In Gombe, we studied two neighboring communities: Kasekela and Mitumba. In Kibale, we studied the chimpanzees of the Kanyawara community.

Gombe is located in western Tanzania, along the shore of Lake Tanganyika (4°40′S, 29°38′E). Gombe has three contiguous communities of chimpanzees, two of which (Kasekela and Mitumba) are well habituated and are followed nearly every day as part of the long-term research at Gombe. During the study period (July 2016-December 2017), the Kasekela community had N = 8 adult males (age ≥ 12 years) and the Mitumba community had N = 5 adult males.

To test if the variation in pant-hoots is explained by geographical differences, we compared calls from Gombe to those of 7 males of the geographically distant Kanyawara chimpanzee community in Kibale (0°33’N, 30°21’E). Following initial observations by Isabirye-Basuta in 1983-1985 (Isabirye-Basuta, 1988), Wrangham initiated the long-term study of Kibale chimpanzees in 1987 (Emery Thompson et al., 2020; Wrangham et al., 1992). We used the calls recorded during October 2010 – September 2011, as a part of another chimpanzee vocal communication study at Kanyawara (Fedurek, Schel, et al., 2013).

### Data collection

N.P.D. and M.L.W. trained two Tanzanian field assistants, Nasibu Zuberi Madumbi and Hashim Issa Salala, to conduct focal follows and record chimpanzee vocalizations at Gombe. They used a Sennheiser ME66 shotgun microphone with K6 power module and a Marantz PMD661 MKII audio recorder. They recorded the vocalizations with a 96 kHz sampling frequency and a 16-bit amplitude resolution. They conducted focal follows of individual males with the goal of recording as many calls as possible from the focal male. In addition to recording calls from the focal target, they also opportunistically recorded as many other calls as possible from known individuals to obtain the maximum number of calls. For each recording, they noted additional information including caller behavior, context, location, and party composition. Here, the recordings were obtained in traveling, feeding, resting, and displaying contexts. However, to ensure sufficient sample sizes and consistency with recordings from Kanyawara, we limited analysis for context differences to calls recorded in traveling and feeding contexts. While the field assistants recorded all call-types from both males and females, here we focus on pant-hoots from males, because (1) pant-hoots have been the focus of previous dialect studies; (2) they can be heard from far away, making them plausible signals of community membership, and (3) males produce pant-hoots more often than females (Wilson et al., 2007).

From July 2016 to December 2017 the team recorded a total of N = 1252 calls (N = 884 from Kasekela and N = 368 from Mitumba). We reviewed these recordings and found that N = 723 (N = 481 from Kasekela and N = 242 from Mitumba) were of sufficiently high quality for acoustic analyses. These recordings consisted of a variety of calls including pant-hoots, pant-grunts, rough-grunts, waa-barks, and screams. Of the pant-hoots in these recordings, some were choruses (where multiple individuals pant-hoot together), and not all were from identified individuals. Choruses that had overlapping elements from multiple callers were excluded, as such overlap makes it harder to extract meaningful acoustic features from known individual callers. Further, to optimize both the number of recordings per individual and the total number of individuals included in the analyses, we excluded individuals that had fewer than 8 pant-hoot call recordings. Based on this criterion, we excluded two individuals: Ferdinand (FE) and Gimli (GIM). While high-ranking males usually call most frequently (Wilson et al., 2007), the highest-ranking male at the start of our study, FE, was overthrown in October 2016, after which we were unable to record any more pant-hoots from him. In Mitumba, in July 2017, the alpha male Edgar (EDG) killed one of the adult males Fansi (FAN) (Massaro et al., 2021). However, we were able to record enough calls from FAN for some of the analyses. With these selection criteria, we identified a total of 214 pant-hoots from known individuals (N = 128 from Kasekela and N = 86 from Mitumba) to be used for the acoustic analysis in this study, with N = 6 adult male chimpanzees from the Kasekela community and N = 5 adult males from the Mitumba community.

In the Kanyawara community, P.F. recorded chimpanzee calls using a Sennheiser ME67 shotgun microphone and a Marantz Professional PMD661 solid-state recorder. He recorded with a 44.1 kHz sampling frequency and a 16-bit amplitude resolution. He obtained the recordings during continuous sampling of focal individuals between October 2010 and September 2011. In addition to recording all calls from the focal individual, he recorded any other vocal interactions between the focal and other individuals in the focal party. Along with the recordings, he noted the identity of the caller who started a vocal bout, the identities of any other callers in a vocal bout, and the context of the vocalizations. Here, the recordings were obtained in traveling and feeding contexts. Using the aforementioned selection criteria, we identified 111 calls from 7 known individuals from Kanyawara to be used in the present study. Table 1 summarizes the number of pant-hoots per individual from three communities.

**Table 1:**
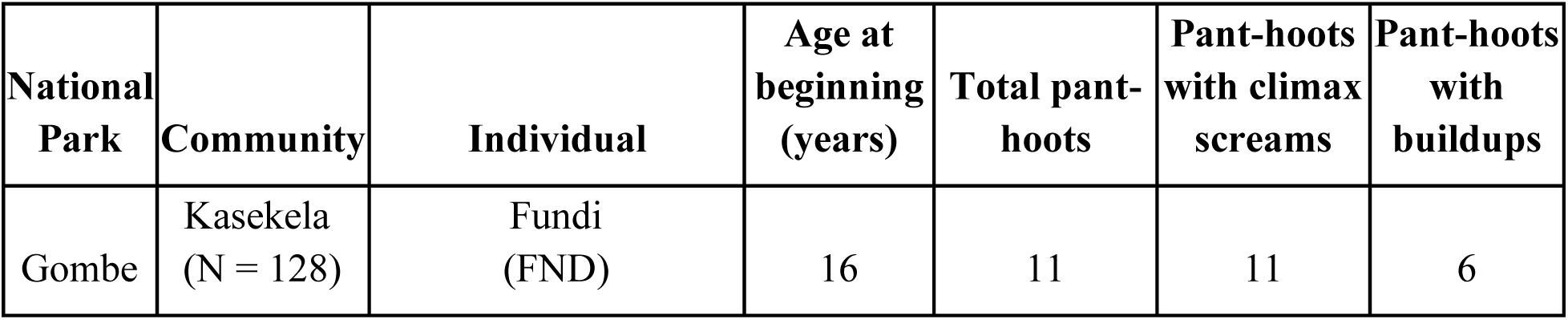
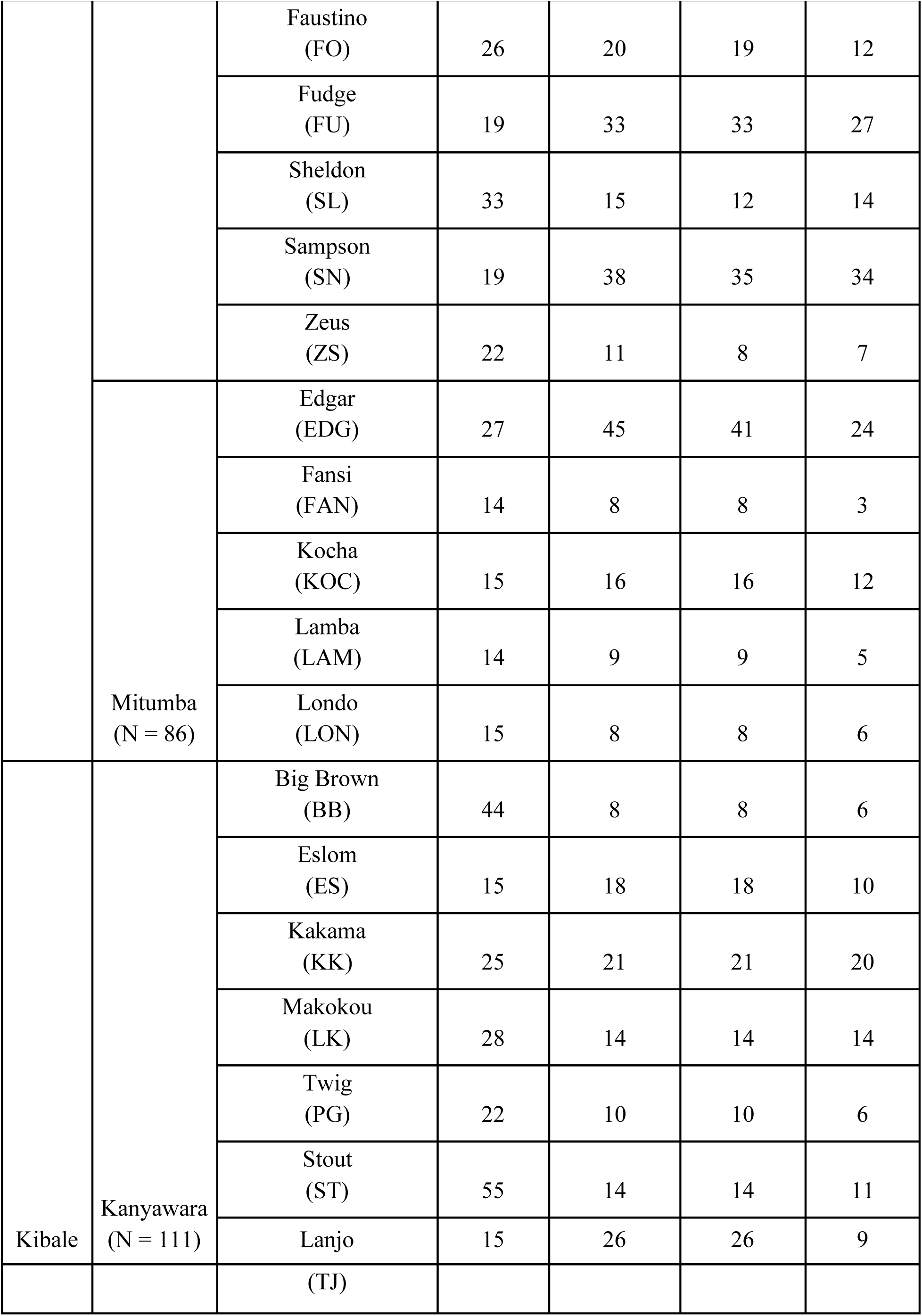
Number of pant-hoots by each individual in the two contiguous Kasekela and Mitumba communities at Gombe National Park, Tanzania and one geographically distant Kanyawara community at Kibale National Park, Uganda included in this study.

### The pant-hoot call

The pant-hoot is a complex call composed of multiple elements (similar to a syllable in human speech). Researchers typically divide pant-hoots into four phases, each of which consists of one or more acoustically similar elements: (i) the introduction, comprised of inhaled and exhaled tonal elements with low fundamental frequencies (generally 300-600 Hz) and a few harmonics, (ii) the build-up, comprised of shorter but more frequent exhaled tonal elements and noisy inhaled elements with low fundamental frequencies (generally 200-500 Hz), (iii) the climax, typically comprised of loud screams of high fundamental frequencies (generally 800-2000 Hz) but which often includes other elements such as hoos and barks, (iv) the letdown, comprised of elements similar to those of the build-up but occurring less frequently and decreasing in fundamental frequency (Figure 1, Sound S1). Chimpanzees do not always produce all four of these phases when giving pant-hoot calls. Sometimes during the call, chimpanzees hit tree buttresses with their feet (and rarely with their hands), producing drum-like sounds (Arcadi & Wallauer, 2013).

**Figure 1:**
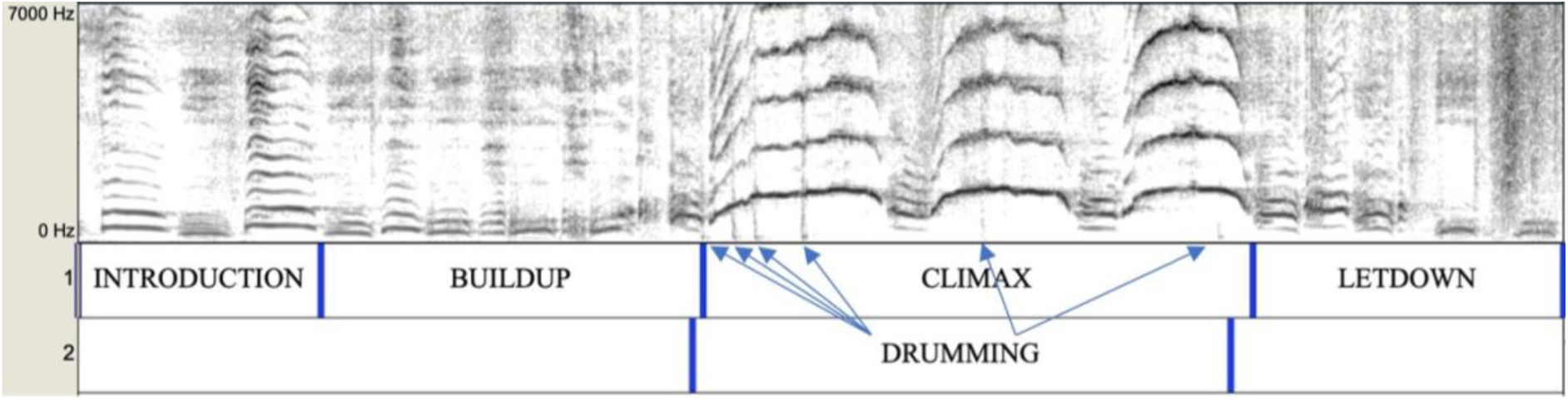
A spectrogram of a typical pant-hoot call with the four phases and drumming labelled.

Differentiating these pant-hoot phases can be difficult as the elements vary substantially in their acoustic structure within each phase. To address this ambiguity and to differentiate systematically among these phases, we proceeded as follows. We identified the exhaled elements in all phases as the elements that reached relatively higher maximum frequencies compared to elements preceding and succeeding them. To distinguish between the introduction and the build-up phase, we defined the start of the build-up as the first exhaled element of markedly shorter duration compared to the previous elements. The buildup consisted of a series of elements with a similarly short duration. Next, to distinguish between the build-up and the climax, we defined the start of the climax as the first exhaled element with a fundamental frequency greater than 500 Hz (see ‘500 Hz’ rule (Mitani et al., 1999)). Next, to distinguish between the climax and the letdown, we defined the end of the climax as the last tonal scream element. In cases where the climax phase did not include screams, we marked the end of the climax as the first element of a reduced fundamental frequency. The letdown phase consisted of a series of these elements of a lower fundamental frequency. Since the different elements of the pant-hoots could often be difficult to distinguish and are subject to observer bias, we only describe the climax in terms of scream and non-scream elements that we found relatively easy to distinguish.

### Acoustic feature extraction

Given the structure of the pant-hoots described above, acoustic analysis of pant-hoots could be performed by measuring acoustic features in different ways. We extracted two main categories of acoustic features from the spectrogram representations of the calls: structural features and spectral features. Structural features describe the composition of the elements in different phases of the pant-hoots and their temporal patterning. For these, we selected 25 acoustic features similar to those used in previous studies of chimpanzee dialects (Crockford et al., 2004; Mitani et al., 1992, 1999) (Table 2). Spectral features quantify the frequency and tonal structure of individual elements from the power spectrum. We measured these from selected specific elements: one build-up element (24 features), and one climax element (25 features). We used semi-automatic measurements of acoustic features using Avisoft-SASLab Pro v. 5.2 (Specht, 2004) and LMA (Fischer et al., 2013; Schrader & Hammerschmidt, 1997) (Table 3).

**Table 2:** Structural acoustic features manually measured using Praat v 6.1.15 that were used in this study. We also indicate which features were used in other studies of chimpanzee dialects. Categorical variables are marked with *. Only numeric variables were used in the multivariate analyses including the PCAs and pDFAs since those techniques do not handle categorical variables. **Drumming related variables were not included in the multivariate analysis due to small sample sizes. However, descriptive plots are included in the supplementary materials.

| **Structural acoustic features used in this study** | **Part of the pant-hoot** | **Crockford et al. 2004** | **Mitani et al. 1999** | **Mitani et al. 1992** |
| --- | --- | --- | --- | --- |
| Duration of the call (from build-up to letdown phases) (s) | Entire call | No | No | No |
| Presence of introduction phase* | Introduction | Yes | No | No |
| Presence of build-up phase* | Build-up | Yes | No | No |
| Number of build-up exhalation elements | Build-up | Yes | No | No |
| Number of build-up elements in the first half of the build-up | Build-up | Yes | No | No |
| Number of build-up elements in the second half of the build-up | Build-up | Yes | No | No |
| Duration of build-up phase (s) | Build-up | Yes | Yes | No |
| Rate of build-up phase (elements/s) | Build-up | Yes | Yes | Yes |
| Rate of first half of build-up phase (elements/s) | Build-up | Yes | No | No |
| Rate of second half of build-up phase (elements/s) | Build-up | Yes | No | No |
| Build-up acceleration (rate of second half - rate of first half) | Build-up | Yes | No | No |

| *Structural acoustic features used in this study* | *Part of the pant-hoot* | *Crockford et al. 2004* | *Mitani et al. 1999* | *Mitani et al. 1992* |
| --- | --- | --- | --- | --- |
| Presence of climax phase* | Climax | Yes | No | No |
| Total number of climax elements (including screams and non-scream elements) | Climax | Yes | No | No |
| Number of screams in climax | Climax | Yes | No | No |
| Proportion of climax elements that are screams | Climax | Yes | No | No |
| Duration of climax phase (s) | Climax | No | No | No |
| Presence of letdown phase* | Letdown | Yes | No | No |
| Number of elements in letdown phase | Letdown | Yes | No | No |
| *Structural acoustic feature(s) NOT used in this study* |  |  |  |  |
| Number of introduction elements | Introduction | Yes | No | No |
| Duration of introduction element | Introduction | No | Yes | No |
| Drumming related features** | Drumming | Yes | No | No |

**Table 3:** Semi-automatically measured acoustic features using LMA from the selected build-up and climax elements compared with other studies. Some acoustic features were not used in this study as they were not measured by the version of LMA available to us.

| *Acoustic feature(s) used in this study* | *Crockford et al. 2004* | *Mitani et al. 1999* | *Mitani et al. 1992* |
| --- | --- | --- | --- |
| Duration of the element (ms) | No (for build-up);<br>Yes (for climax) | Yes | Yes |
| Start, end, maximum, minimum, and mean fundamental frequency F0 (Hz) | No (for build-up);<br>Yes (for climax) | No (start and end);<br>Yes (maximum, minimum, and mean) | No |
| Frequency range of F0 (Maximum F0 - Minimum F0) (Hz) | No | Yes | Yes |
| Tonality measures: mean and maximum frequency difference between the original F0 curve and the floating average curve (Hz) | No (for build-up);<br>Yes (for climax) | No | No |
| Location of maximum F0 relative to the duration ([1/duration]*location) | No (for build-up);<br>Yes (for climax) | No | No |
| Factor of linear trend of F0 (measures if the F0 is rising, falling, or flat on average) | No (for build-up);<br>Yes (for climax) | No | No |
| Mean and maximum deviations between F0 and linear trend line (Hz) | No (for build-up);<br>Yes (for climax) | No | No |
| Start, end, maximum, minimum, and mean peak frequencies (Hz) | No (for build-up);<br>Yes (for climax) | No | No |
| Peak frequencies with maximum and minimum amplitude (Hz) | No (for build-up);<br>Yes (for climax) | No | No |
| Locations of maximum and minimum peak frequencies relative to the duration ([1/duration]*location) | No | No | No |
| Maximum difference between peak frequency values in successive time segments (Hz) | No (for build-up);<br>Yes (for climax) | No | No |
| Mean and maximum wiener entropy coefficient (0-1; 1=noise) | No | No | No |
| *Acoustic feature(s) NOT used in this study* |  |  |  |
| Slope of F0 from start to maximum (Hz/ms) | Yes | No | No |
| Slope of peak frequency from start to maximum (Hz/ms) | Yes | No | No |
| Maximum F0 start F0 (Hz) and Maximum F0 minimum F0 (Hz) | Yes | No | No |
| F0 at midpoint of introduction element (Hz) | Yes | Yes | No |
| Peak frequency at midpoint of inhaled elements (Hz) | Yes | No | No |
| Peak frequency at midpoint of exhaled elements (Hz) | Yes | No | No |
| Peak frequency of inhaled - peak frequency of exhaled elements (Hz) | Yes | No | No |
| Ratio of F1/F2, the first and second formant frequencies | No | Yes | No |
| Bandwidth (Hz) | No | Yes | No |

We extracted the acoustic features as follows. First, we measured structural features from the pant-hoot phases by visually inspecting spectrograms of entire pant-hoots using Praat version 6.1.15. We considered each phase separately and measured a set of acoustic features from each phase (Table 2). We present the visual summaries of these structural features of the pant-hoots from the three communities in the supplementary materials (Figures S1 (a-m)). Next, for the semi-automatic extraction of acoustic features, we chose one element from the buildup phase and one element from the climax phase. From the build-up phase, we chose the middle element in case of an odd number of build-up elements, and the element immediately preceding the middle of the build-up in case of an even number of elements (Mitani et al., 1999). From the climax phase, we chose the scream that reached the highest fundamental frequency in the spectrogram. To obtain appropriate frequency and time resolutions, we down sampled the sampling frequency to 24 kHz using Avisoft-SASLab Pro, resulting in a frequency range of 12 kHz. Next, using Avisoft-SASLab Pro, we created spectrograms with an FFT length of 1024 points, frame size of 100%, and Hamming window with an overlap of 93.75%. This resulted in a frequency resolution of 23 Hz, and a time resolution of 2.7 ms, which is sufficient to reveal tonal properties and extract acoustic features from buildup and climax elements. We then imported the spectrograms in LMA and extracted acoustic features (listed in Table 3) using the harmonic cursor tool. We did not extract additional acoustic features from elements in the introduction and the letdown phases as the introduction was not always recorded fully, and the letdown exhibited high variability in the type of elements, making comparison among letdowns difficult. We attempted to be consistent with previous studies by including as many acoustic features used in previous studies as possible (Tables 2, 3). However, some acoustic features used in previous studies could not be measured using the software packages available to us. Nevertheless, we consider that the acoustic features we used should encompass the relevant range of variation in chimpanzee pant-hoots, without loss of generality.

### Statistical analysis

To confidently make conclusions about whether chimpanzees have community-specific dialects that are an outcome of vocal learning, we need to control for confounding factors. If chimpanzees learn community-specific calls to distinguish themselves from members of neighboring communities, then calls from neighbors should be more distinct than calls from distant communities. Hence, we first need to ensure that the community differences (if any) are not merely differences that could be attributed to geographically distant communities. Geographically distant communities could also have other confounding genetic and environmental differences that can lead to differences in vocalizations, but not because of vocal learning (Mitani et al., 1999). Secondly, we need to control for the context of vocalization as pant-hoots given in different contexts may be acoustically different (Notman & Rendall, 2005; Uhlenbroek, 1996). And lastly, we need to control for the individual variation as having multiple calls from the same individual violates the assumption of independence, and can lead to erroneous inference about group differences (Mundry & Sommer, 2007). To address these, we performed the analyses as follows.

We used the permuted Discriminant Functions Analysis (pDFA) procedure to test for differences in the factor of interest (a.k.a. test factor) while controlling for a confounding factor (a.k.a. control factor) (Mundry & Sommer, 2007). The traditional Discriminant Functions Analysis (DFA) technique handles only one factor at a time and is known to inflate group differences in two-factorial designs with a confounding factor (individual identity is among the most common confounding factors in similar studies (*ibid.*)). The pDFA procedure allows two-factor designs by performing a permutation test on the classification accuracy of DFA. The permutation test tests if the observed classification accuracy of DFA is significantly higher than expected by chance while accounting for the accuracy inflating effect of a confounding factor. The procedure works as follows. Firstly, the procedure samples a specified number of balanced and randomized datasets from the original dataset. It randomizes the labels of the test factor based on the combinations of different categories of test and control factors. It performs the balancing such that there is the same number of observations from each category of the test factor. Next, it performs a traditional DFA on each of these randomized datasets and obtains a distribution of classification accuracies of these DFAs. The observations left out due to balancing are used for cross-validation to obtain the out-of-sample, cross-validated classification accuracies. The distribution of classification accuracies of randomized datasets describes the probabilities of obtaining particular classification accuracies using a traditional DFA just based on chance. The expected value from this distribution is compared with the classification accuracy obtained on the original dataset (observed classification accuracy) to obtain a p-value for the permutation test. In other words, this distribution provides an estimate of the inflation in classification accuracy of the test factor caused by the confounding effect of the control factor.

We performed 1000 permutations (*i.e.*, 1000 randomized datasets including the original dataset) in each of our analyses and used an alpha level of **α** = 0.05 on the cross-validated classification accuracy to infer a significant difference. The test factors of interest in our study were context, community identity, and individual identity. And the control factors were context and individual identity, depending on the analysis. For different test factors in the pDFA, we had to consider the data designs to ensure proper randomization and balancing of the permuted datasets. In our study, two design situations occurred: crossed and nested. A crossed design occurs when all the categories of the test factor are recorded in all categories of the control factor. So, a pDFA with crossed design could only be used in testing for differences in context by only including individuals recorded in both contexts. While testing for differences in community identity and individual identity, a nested design occurs. In a nested design, the categories of the control factor are nested within categories of the test factor, or the categories of the test factor are nested within some other factor known as the restriction factor. Since individual identity is nested within community identity, a nested design occurs when testing for differences in the community or individual identities.

We performed all statistical analyses in R version 4.0.2 using RStudio version 1.3.1093. The pDFAs were carried out using a set to R functions provided by R. Mundry. These functions implement the pDFA procedure and are built on top of the *lda* function in the R package MASS (Ripley et al., 2020) that is used to perform traditional DFAs. We performed four different analyses for different kinds of acoustic features: (i) structural features (Table 2), (ii) build-up element features (Table 3), (iii) climax scream features (Table 3), and (iv) all features combined (Tables 2 and 3). We tested for differences in context, community identity, and individual identity in each of these four types of acoustic features. For each kind of acoustic feature set, we tested for context before performing other analyses to determine whether context was a significant confounding factor that needed to be statistically controlled for in the subsequent analyses. To control for geographical differences, we performed two separate analyses for each kind of acoustic feature set. Following Crockford et al. (2004), we first investigated the acoustic structure of pant hoots from the two neighboring communities of Gombe, where maximal differences were expected. In order to then compare the two Gombe communities to a geographically distant community, we ran pDFAs including all three communities. To control for individual differences, we used individual identity as the control factor in each of the pDFAs when testing for context and community identity. When testing for individual differences using individual identity as the test factor, we used community identity as the restriction factor. When a restriction factor is added in the pDFA, the randomization process is done while accounting for the fact that the test factor is nested within the restriction factor. To avoid overfitting, we ensured that only as many, or fewer acoustic features are used to perform the DFAs as there are observations in the category of the test factor with the fewest observations. In cases when there were more acoustic features, we used Principal Components Analysis (PCA) on the acoustic features to reduce the number of features while accounting for most variation contained in different acoustic features. We used the scores of each observation on the principal components as the features to be used in the DFAs. We used as many principal components as there were number of observations in that category of the test factor with the fewest observations or as many principal components that explained 90% of the variation, whichever was smaller. However, to ensure that the large number of variables was not leading to overfitting, we verified the consistency of the results of the pDFAs over different numbers of selected principal components using Cattell’s scree test (Cattell, 1966).

For each acoustic feature set used in the analyses, we chose a subset of recordings based on the following criteria. For structural features, we first removed acoustic features that had too many missing values for sufficient statistical power. These mainly included acoustic features related to drumming, as only 18% of the recorded pant-hoots had drumming (Table 2, Figures S1 (l-m)).

After that, we removed categorical features that indicated the presence or absence of the four phases as categorical features are not handled by DFAs. Next, we removed the cases that had missing values in any of the remaining 14 acoustic features. While the categorical features were eliminated, the information contained in them was included in other features that indicated the number of elements in each phase. A value of 0 in those features would indicate absence of a phase, whereas a non-zero value would indicate the presence. For the build-up feature set, we only included pant-hoots that included the build-up phase. Similarly, for the climax feature set, we only included pant-hoots that included the climax phase. And lastly, while including all the acoustic features simultaneously (structural, build-up, and climax features), we only included pant-hoots that had both buildup and climax phases.

In case a pDFA revealed significant differences, we used the *repDFA* function written by C. Neumann (Berthet et al., 2017; Neumann, 2020) to identify the key variables that allow discrimination of the test factor. This function creates 1000 balanced datasets in crossed designs, re-runs 1000 DFAs and records the variable that had the highest coefficient on the first and second linear discriminant functions in each of those DFAs. Variables that have the highest coefficient in many of those permutations are arguably the most important in discriminating the test factor. For nested designs, we modified Neumann’s function and wrote a new function called *repDFA_nested* (available on GitHub). This function modified *repDFA* such that it created balanced datasets by randomly sampling with replacement the same number of recordings for each individual in the analysis. We further tested the significance of the individual variables identified with *repDFA* and *repDFA_nested* using Generalized Linear Mixed Models (GLMMs). We controlled for individual identity in the GLMMs by including individual IDs as random intercepts and adjusted the p-values for multiple comparisons using Benjamini–Hochberg adjustment method.

### Data sharing statement

The R code and data for the analyses are available from GitHub at https://github.com/desai-nisarg/Gombe-dialects. Audio recordings from Gombe are available from M.L.W. and from Kanyawara available from P.F. at reasonable request.

### Ethical note

The research reported in this paper is based on data collected non-invasively from free-ranging chimpanzees. The Institutional Animal Care and Use Committee at the University of Minnesota did not require a review due to the purely observational nature of the research. Research at Gombe National Park was performed with approval from the Tanzania Wildlife Research Institute and the Tanzania Commission for Science and Technology and adhered to additional ethical guidelines set by the Jane Goodall Institute.

Research at Kibale National Park was approved by the Department of Psychology Ethics panel at the University of York and permission to conduct the study was granted by the Ugandan Wildlife Authority and the Ugandan National Council for Science and Technology. The study complied with the laws of Uganda.

## Results

### Differences in pant-hoots between contexts

We found a statistically significant difference in the structural acoustic features (Table 2) between feeding and traveling contexts after controlling for individual identity (pDFA with structural features, observed classification accuracy: 63.9 % vs. expected by chance: 51.7 %; p = 0.044, Table 4). Using the *repDFA* function, we identified principal component 6 (PC6) to be the best discriminator of the contexts in 716 out of the 1000 DFAs followed by principal component 1 (PC1) in 191 out of the 1000 DFAs. PC6 loaded most heavily on the number of letdown elements and buildup acceleration had the second highest loading. PC1 loaded most heavily on the number of buildup elements but other features of the buildup had comparable loadings. These features were: duration of the buildup phase, rate of the buildup phase, number of elements in the first and second half of the buildup, and rate of the first and the second half of the buildup. Further, the principal components plot made by performing principal components analysis on the structural features shows a distinct band of calls given mostly in feeding contexts (Figure 2(a)). Principal component 2 (PC2) explained the most variance in this band and it loaded most heavily on the number of climax elements. These were pant-hoots with a greater than average number of climax elements, and which did not have a buildup phase. We performed significance tests for the highest loading features of these three important components using Poisson GLMMs and controlled for individual identity by including it as a random effect in the models. All three of these acoustic features were statistically significantly different between contexts. (i) Number of letdown elements, β (Travel) = 0.57, Benjamini–Hochberg adjusted p-value = 3.8e-07. (ii) Number of buildup elements, β (Travel) = 0.34, Benjamini–Hochberg adjusted p-value = 4.8e-06. (iii) Number of climax elements, β (Travel) = -0.27, Benjamini–Hochberg adjusted p-value = 1.8e-03 (Figures S2(a-c)). Hence, we conclude that the number of elements in the build-up, the climax, and the letdown phases potentially encoded contextual information.

**Figure 2:**
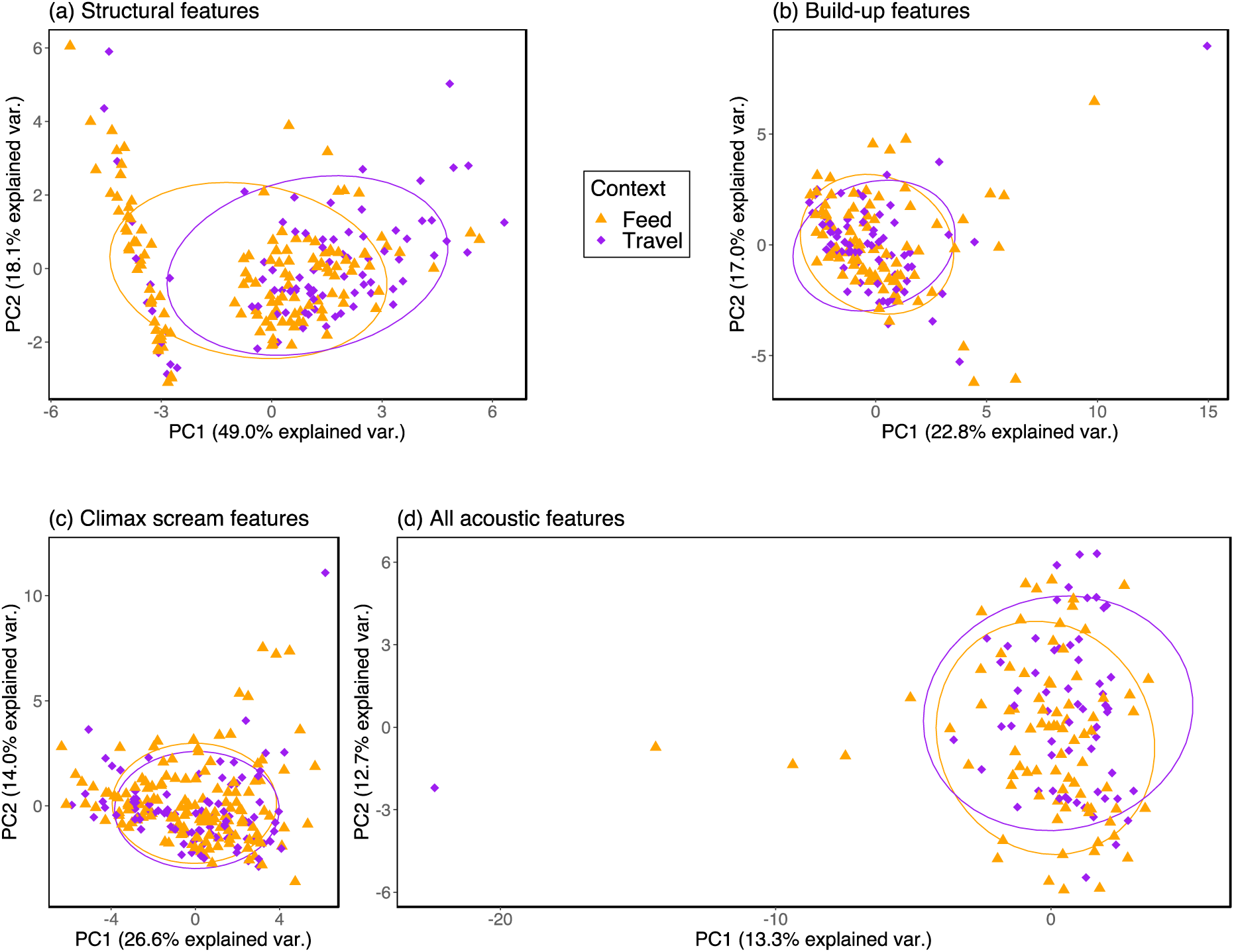
Principal components plots with the 68% normal data ellipses containing 68% of the data points included for each context. (a) Principal Components Analysis performed on structural features. Pant-hoots given in different context separate over PC1. (b) Principal Components Analysis performed on acoustic features of the selected build-up element. (c) Principal Components Analysis performed on acoustic features of the selected climax element. (d) Principal Components Analysis performed on all acoustic features simultaneously from all three communities. (b), (c), and (d) reveal strong overlap between contexts.

**Table 4:**
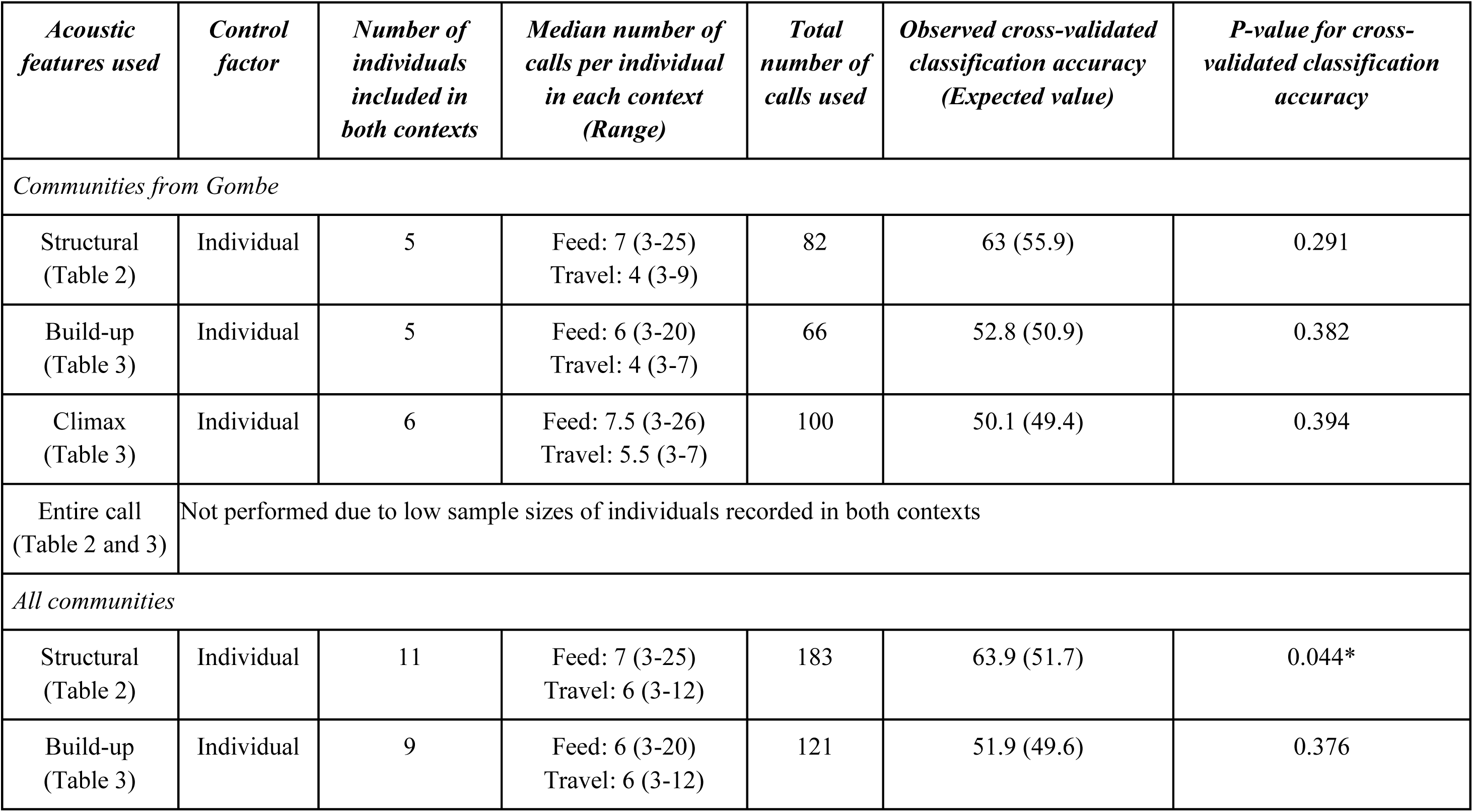
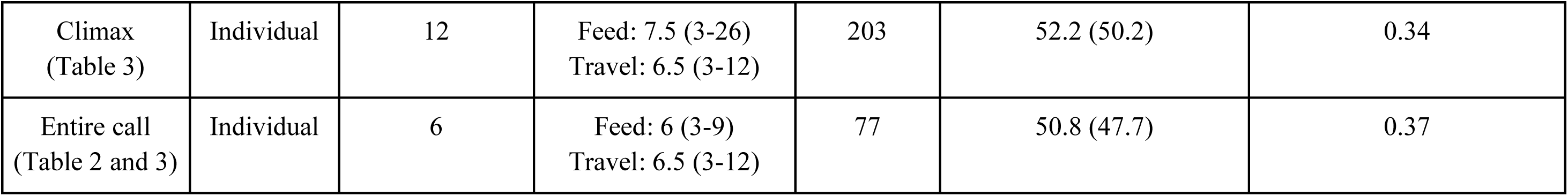
Summary of the results from the pDFAs with context as the test factor and individual identity as the control factor for different types of acoustic features. We indicate the number of individuals recorded in both feeding and traveling contexts, the range of number of calls per individual and the total number of calls considered for each of the analyses.

Further, we found no differences in the contexts in other types of acoustic features after controlling for individual identity among the contiguous communities or all communities taken together. Cross-validated p-values for pDFA performed on all communities with (i) build-ups: p = 0.38, (ii) climax screams: p = 0.34, and (iii) all acoustic features simultaneously: p = 0.37. Cross-validated p-values for pDFA performed on communities within Gombe with (i) structural features: p = 0.29, (ii) build-ups: p = 0.38, and (iii) climax screams: p = 0.39 (Table 4). Figures 2(b-d) show the overlap between contexts in these acoustic features in a multidimensional space. Hence, we controlled for context when testing for differences in structural features alone and not when testing for differences in other types of acoustic features in the subsequent analyses.

### Differences in pant-hoots among communities of chimpanzees

Figures 3(a-d) show the clusters of the three communities in multidimensional spaces of the structural features, build-ups, climaxes, and all features taken simultaneously. These show strong overlap among communities, suggesting a lack of community-level differences that is confirmed by the pDFAs, with one exception: a statistically significant difference in acoustic features of the climax scream among the communities when we included the geographically distant Kanyawara community in the analysis (pDFA on climaxes of all communities, observed classification accuracy: 54 % vs. expected: 40.8 %; p = 0.016). These features did not differ statistically between the two neighboring communities at Gombe with alpha = 0.05 but would differ with alpha = 0.10 (observed classification accuracy: 70.7 % vs. expected: 58 %; p = 0.089). The *repDFA_nested* function identified principal component 4 (PC4) and principal component 2 (PC2) to be the best discriminators of community identity for the first two discriminant functions of climax features. The top three variables with the largest loadings on the principal component 4 (PC4) of the climax scream features were factor of linear trend of F0, duration of climax scream, and the location of maximum F0 relative to duration. The top three variables with the largest loadings on the principal component 2 (PC2) of the climax scream features were maximum Wiener entropy coefficient, mean Wiener entropy coefficient, and maximum difference in peak frequency between successive time segments. We performed significance tests on these features using Linear Mixed Models and controlled for individual identity by including it as a random effect in the models. Features on the PC4 were not significantly different among communities. (i) Factor of linear trend of F0, Benjamini–Hochberg adjusted p-value = 0.83 (Kanyawara - Kasekela), 0.078 (Kanyawara - Mitumba). (ii) Duration of climax scream, Benjamini–Hochberg adjusted p-value = 0.72 (Kanyawara - Kasekela), 0.27 (Kanyawara - Mitumba). (iii) The location of maximum F0 relative to duration, Benjamini–Hochberg adjusted p-value = 0.72 (Kanyawara - Kasekela), 0.15 (Kanyawara - Mitumba). Features on the PC2 were significantly different among communities. (iv) Mean Wiener entropy coefficient, Benjamini–Hochberg adjusted p-value = 0.011 (Kanyawara - Kasekela), p = 3.5e-06 (Kanyawara - Mitumba). (v) Maximum Wiener entropy coefficient, Benjamini–Hochberg adjusted p-value = 1.4e-05 (Kanyawara - Kasekela), p = 3.2e-07 (Kanyawara - Mitumba). (vi) Maximum difference in peak frequency between successive time segments, Benjamini–Hochberg adjusted p-value = 6.5e-04 (Kanyawara - Kasekela), p = 0.72 (Kanyawara - Mitumba).

**Figure 3:**
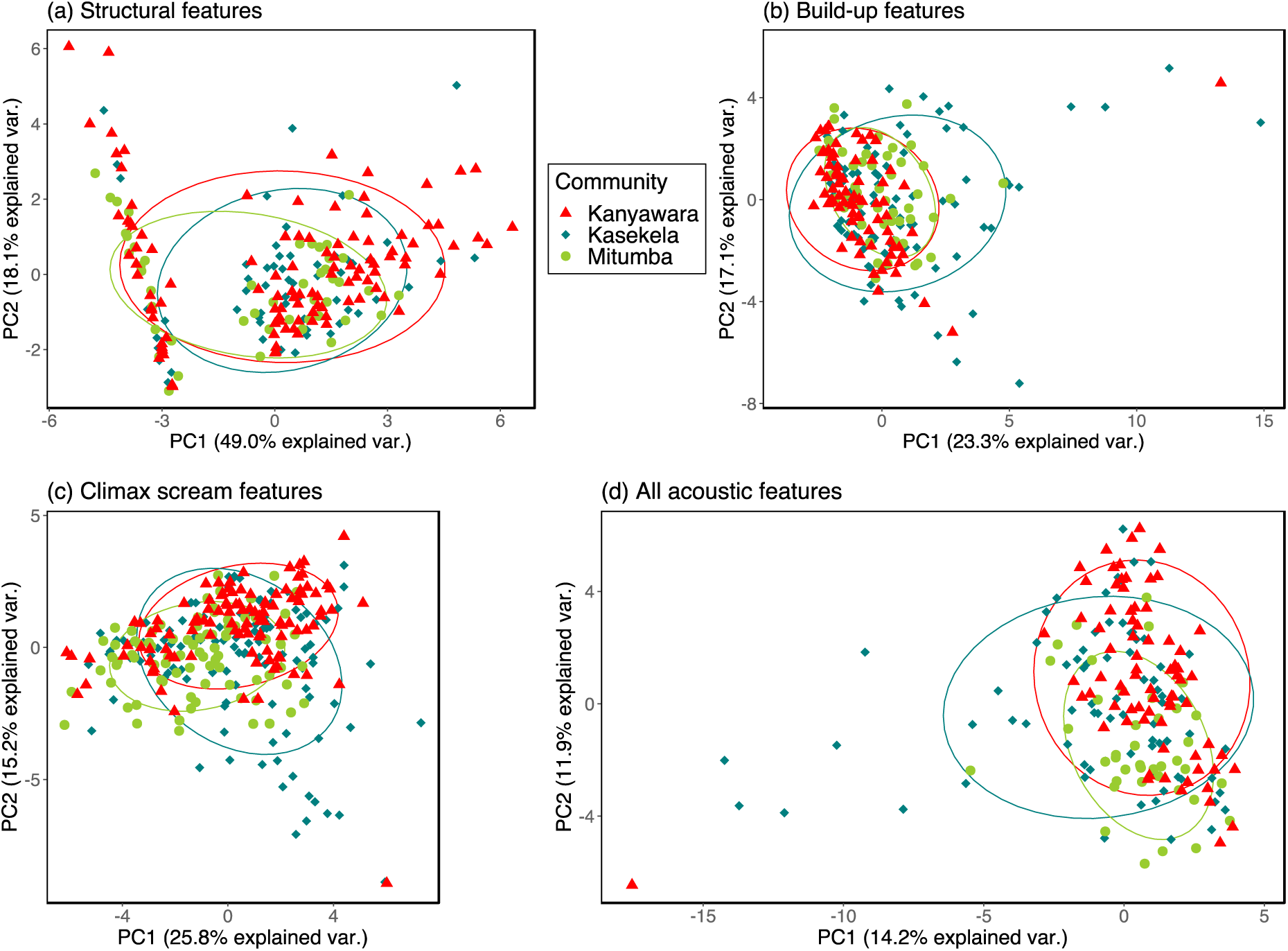
Principal Components plots with the 68% normal data ellipses containing 68% of the data points included for each community. (a) Principal Components Analysis performed on structural features. (b) Principal Components Analysis performed on acoustic features of the selected build-up element. (c) Principal Components Analysis performed on acoustic features of the selected climax element. Kasekela and geographically distant Kanyawara communities separate to some extent over PC2. (d) Principal Components Analysis on all acoustic features simultaneously from all three communities. (a), (b), and (d) reveal strong overlap among communities.

Additionally, we observed no differences among the contiguous communities, or all communities taken together in the structural features (controlled for individual identity or context), acoustic features of the build-ups, or all acoustic features considered together (Table 5).

**Table 5:**
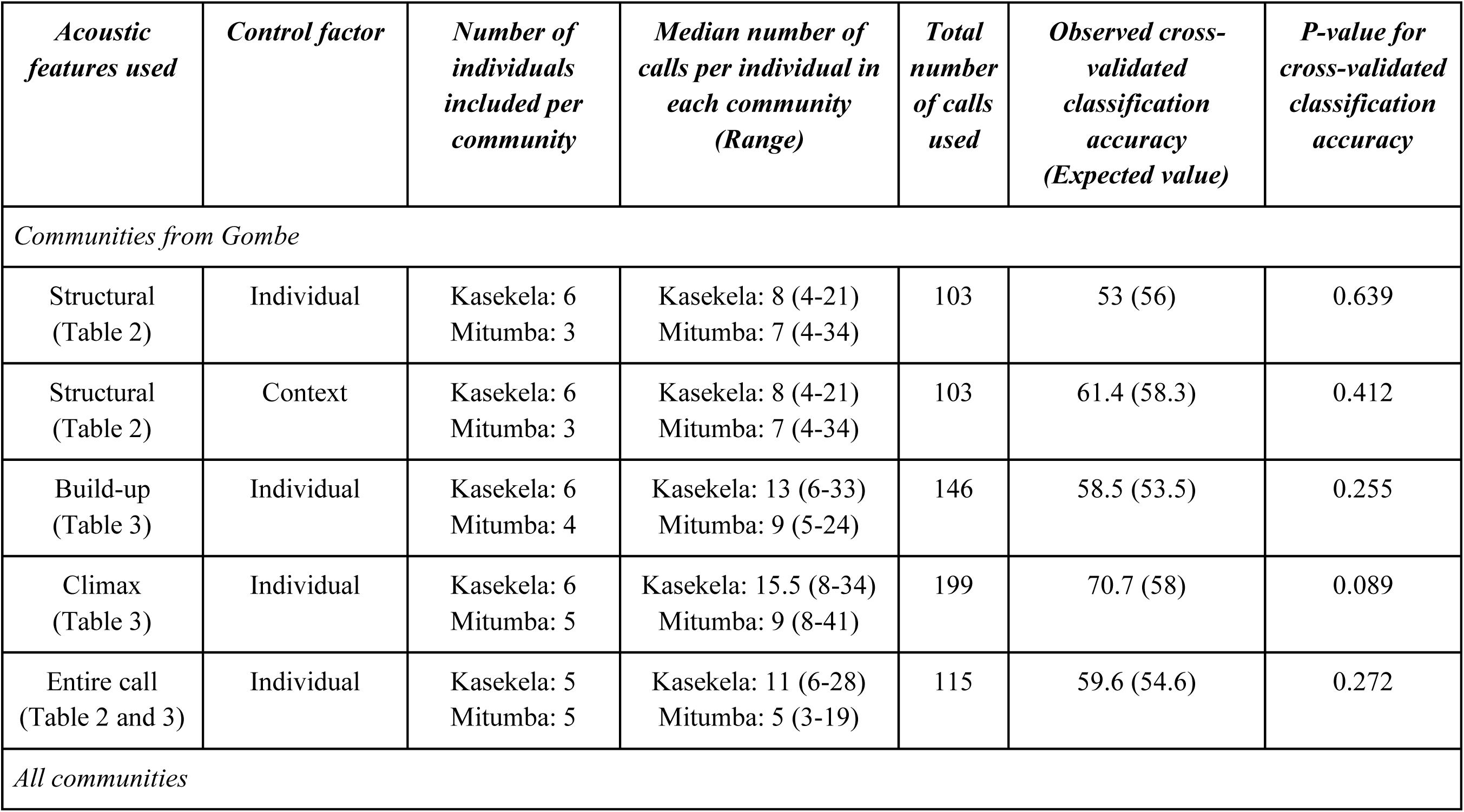
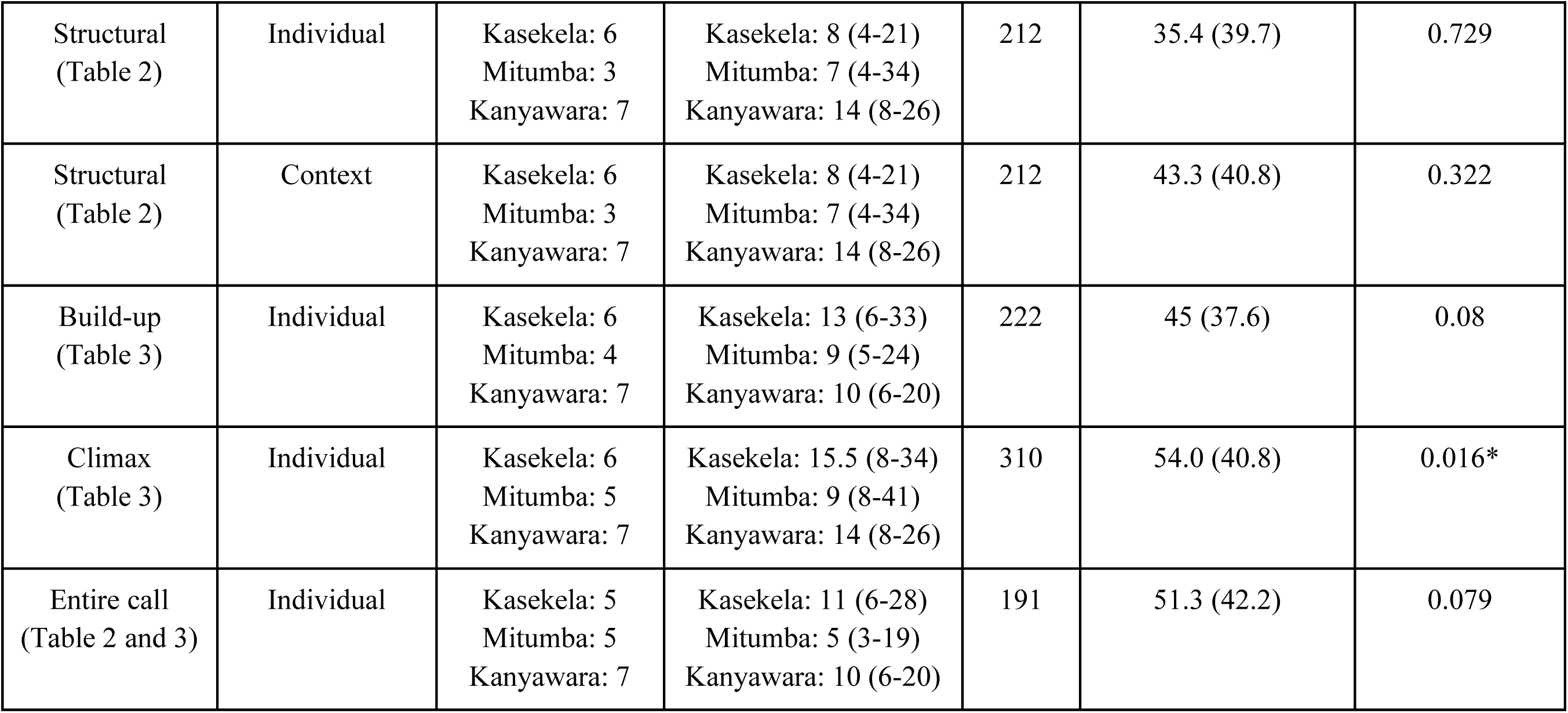
Summary of the results from the pDFAs with community identity as the test factor for different types of acoustic features. We indicate the control factor, the number of individuals from each community, the range of number of calls per individual and the total number of calls considered for each of the analyses. We used context as a control factor only in case of structural features since there was a difference between contexts only in structural features.

### Differences in pant-hoots among individuals

We observed statistically significant differences among the individuals in the structural features, acoustic features of the climax screams, and all acoustic features taken simultaneously. This was true when all communities were taken together as well as when the geographically adjacent communities of Gombe were assessed separately (Table 6). However, the individuals could not be separated based on acoustic features of the selected build-up elements in any setting (pDFA on build-up features of all communities: p = 0.18, and Gombe: p = 0.15; Table 6).

**Table 6:** Summary of the results from the pDFAs with individual identity as the test factor for different types of acoustic features. We used community ID as the restriction factor except when using context as a control factor. We indicate the number of individuals included, the range of number of calls per individual and the total number of calls considered for each of the analyses.

| *Acoustic features used* | *Control or restriction factor* | *Number of individuals included* | *Median number of calls per individual (Range)* | *Total number of calls used* | *Observed cross-validated classification accuracy (Expected value)* | *P-value for cross-validated classification accuracy* |
| --- | --- | --- | --- | --- | --- | --- |
| *Communities from Gombe* |  |  |  |  |  |  |
| Structural (Table 2) | Community (restriction factor) | 11 | 13 (4-41) | 171 | 24 (10.3) | 0.001* |
| Structural (Table 2) | Not performed due to low sample sizes of individuals recorded in both contexts |  |  |  |  |  |
| Build-up (Table 3) | Community (restriction factor) | 10 | 12 (5-33) | 146 | 16.1 (12.1) | 0.15 |
| Climax (Table 3) | Community (restriction factor) | 11 | 12 (8-41) | 199 | 24.2 (13.2) | 0.006* |
| Entire call<br>(Table 2 and 3) | Community<br>(restriction<br>factor) | 10 | 10 (3-28) | 115 | 23.5 (13.2) | 0.024* |
| *All communities* |  |  |  |  |  |  |
| Structural<br>(Table 2) | Community<br>(restriction<br>factor) | 18 | 13.5 (4-41) | 280 | 19.5 (6.9) | 0.001* |
| Structural<br>(Table 2) | Context<br>(control factor) | 7 | 14 (9-34) | 119 | 35.8 (24.7) | 0.043* |
| Build-up<br>(Table 3) | Community<br>(restriction<br>factor) | 17 | 11 (5-33) | 222 | 10.6 (8.2) | 0.18 |
| Climax<br>(Table 3) | Community<br>(restriction<br>factor) | 18 | 14 (8-41) | 310 | 20.1 (9.4) | 0.001* |
| Entire call<br>(Table 2 and 3) | Community<br>(restriction<br>factor) | 17 | 10 (3-28) | 191 | 14.4 (7.6) | 0.007* |

Figures 4(a-c) show the differences among individuals of the three communities in the multidimensional space all acoustic features taken simultaneously. In Kasekela, calls from the individuals FND, FU and the pair FO and SL separate over PC2. While calls from FO, SL, and SN overlap, SN could be differentiated to some extent on PC1 (Figure 4(a)). In Mitumba, while calls from the individuals EDG and LAM overlap, they could be differentiated from KOC, LON, and FAN from a combination of PC1 and PC2 values (Figure 4(b)). In Kanyawara, calls from the individuals BB, ES, and TJ separate from LK on PC1 and from PG and KK on PC2 (Figure 4(c)).

**Figure 4:**
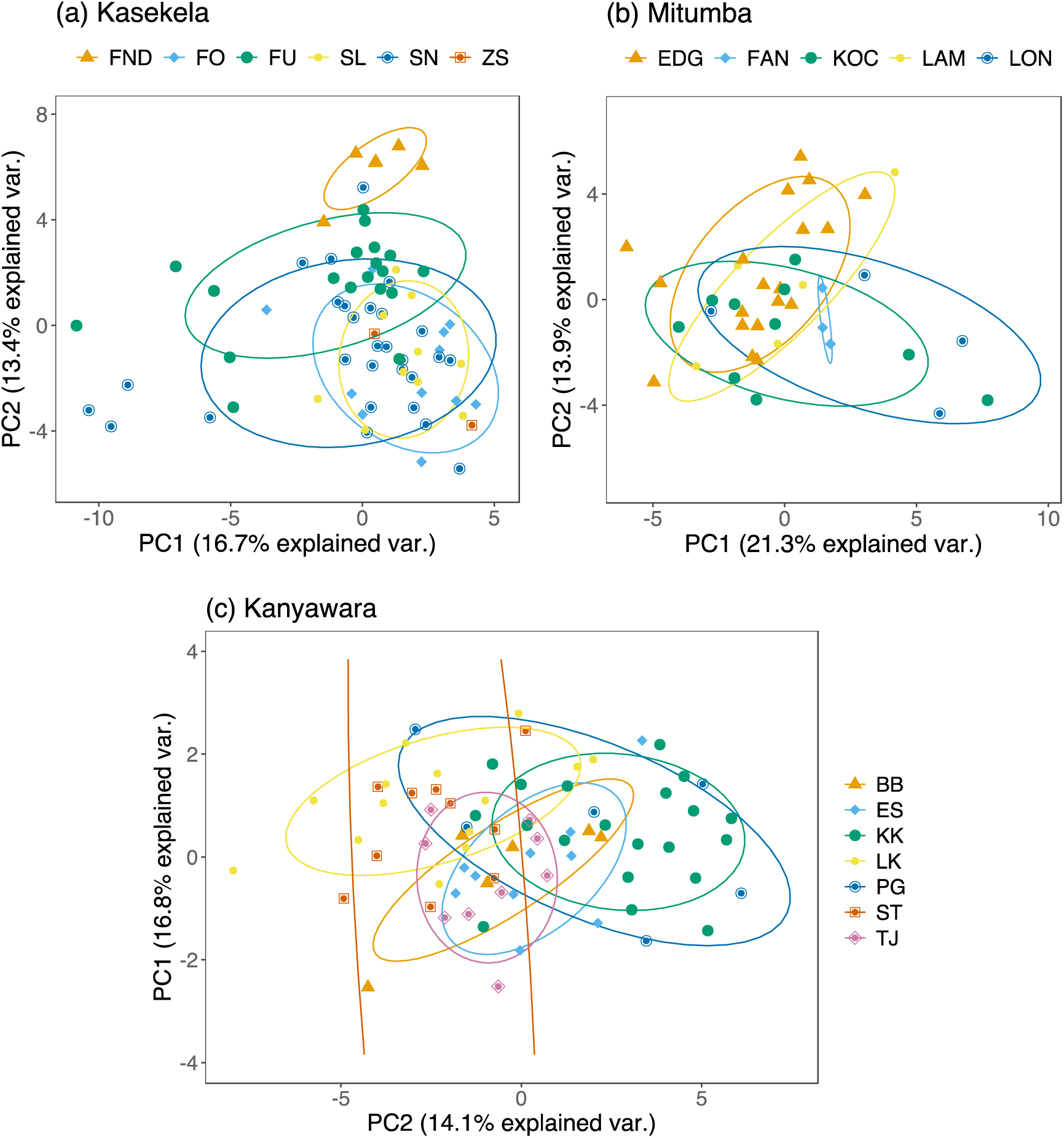
Principal Components plots with the 68% normal data ellipses containing 68% of the data points included for each individual. Principal Components Analysis was performed on the structural features as well as features of the selected build-up and climax elements simultaneously from the three communities. The 68% normal data ellipses revealed a lower overlap compared to community identity and context. Plot for (a) Kasekala. Some individuals formed distinct clusters over PC2. (b) Mitumba. Some individuals formed distinct clusters over a combination of PC1 and PC2. (c) Kanyawara. Some individuals formed distinct clusters over PC2 and others over PC1.

For the structural features, the *repDFA_nested* function identified principal component 2 (PC2) and principal component 7 (PC7) to have the highest loadings on both discriminant function 1 (PC2 higher than PC7) and discriminant function 2 (PC7 higher than PC2). Combined they had the highest loadings on discriminant function 1 in 619 out of 1000 DFAs and on discriminant function 2 in 423 out of 1000 DFAs. Top three acoustic features with the highest loadings on PC2 were the number of climax elements, build-up to letdown duration, and the duration of climax. And on PC7, the number of climax screams, build-up to letdown duration, and the duration of climax loaded the highest. For the climax features, the *repDFA_nested* function identified principal component 1 (PC1) to have the highest loading on discriminant function 1 in 860 out of 1000 DFAs and principal component 3 (PC3) to have the highest loading on discriminant function 2 in 842 out of 1000 DFAs. Top three acoustic features with the highest loadings on PC1 were mean F0, maximum F0, and frequency range of F0. And on PC3 were minimum F0, minimum peak frequency, and start frequency of F0. Lastly, when all features were taken together, PC2 loaded the highest on 945 out of 1000 DFAs. And the top three acoustic features that loaded the highest on PC2 were maximum F0, mean F0, and frequency range of F0. This confirmed the findings above and also suggested that the strongest individual signal was in the acoustic features of the fundamental frequency in the climax scream. We do not report the results from the GLMMs for these acoustic features as differences among specific individuals are not of general interest. However, we observed statistically significant differences in some (but not all) pairs of individuals in each of these acoustic features suggesting that some individuals could be identified with more certainty than others. We can see this reflected in the low classification accuracies in the pDFAs (Table 6).

## Discussion

Our analysis of multiple acoustic features of chimpanzee pant-hoots found that pant-hoots could not be distinguished reliably based on the community identity, but instead reflected individual identity. The pant-hoots differed among the communities in only one type of acoustic features (the acoustic features of the climax scream), and only when we included the geographically distant Kanyawara community in the analysis. We did not observe a statistically significant difference in the climax screams of the geographically adjacent communities of Gombe. We also did not observe any differences among the communities in either the structural features, the build-up features, or when taking all the acoustic features simultaneously. The pant-hoots differed most substantially among individuals, irrespective of the inclusion of the geographically distant community in our analyses. The acoustic features of the climax scream element and the structural acoustic features distinguished the individuals, whereas the acoustic features from the build-up element alone did not. Our findings indicate that individual differences are more prominent than group differences in the acoustic structure of chimpanzee pant-hoots. We thus found support for our second hypothesis and not the first hypothesis.

We found that the context of the vocalization could be identified from some structural acoustic features but not from any other kind of acoustic features. Within the structural features, the number of climax elements was higher in feeding contexts and the number of letdown elements as well as buildup elements was higher in traveling contexts. These results are generally consistent with several previous studies. Our results support the findings of Clark & Wrangham (1993), and Fedurek et al. (2016) in finding an association of the letdown phase with the context of the pant-hoot. However, we did not have sufficiently detailed behavioral data to distinguish food arrival pant-hoots separately, and hence, we could not confirm Clark & Wrangham’s finding, that a higher proportion of pant-hoots with letdowns occurred in the context of arrival at a food source. We further observed two more differences that have not been reported previously. First, we found an association of pant-hoots given in feeding contexts with the number of climax elements. Second, we observed a higher number of buildup elements in travel context. Furthermore, we found no differences between the contexts in other acoustic features that describe the tonal properties of the build-up and climax elements. Uhlenbroek (1996) described different types of pant-hoots based on their tonal and spectral properties. She called a tonal pant-hoot with clear harmonic structure and a power spectrum with clear peaks a ‘wail-like’ pant-hoot, and a more noisy pant-hoot lacking a clear harmonic structure and a more evenly distributed power spectrum as a ‘roar-like’ pant-hoot. Uhlenbroek (1996) and Notman & Rendall (2005) found that pant-hoots given in traveling contexts were more ‘roar-like’ and those given in feeding contexts were more ‘wail-like.’ Since we found no context differences in the acoustic features related to the tonal properties, fundamental frequency, noise, or peak frequency, we could not confirm the findings from Uhlenbroek (1996) or Notman & Rendall (2005). Our results indicate that a more fine-grained differentiation of contexts while recording pant-hoots may be needed to distinguish arrival pant-hoots as well as pant-hoots from other contexts such as resting, grooming, displaying. Additionally, our findings suggest that future studies should pay special attention to the structural features whenever the context of pant-hoot production is relevant to the analysis.

We did not find evidence of community-specific differences that would suggest vocal learning. This is contrary to previous studies looking at community-specific differences (Crockford et al., 2004; Marshall et al., 1999; Mitani et al., 1992). In the first study reporting vocal dialects in chimpanzees, Mitani et al. (1992) found differences in geographically distant communities of Gombe and Mahale National Parks and Mahale and Kibale National Parks respectively. We found less evidence of acoustic differences among distant chimpanzees: only in some of the acoustic features of the climax element and not in the build-up element. Among the specific acoustic features of the climax element, Mitani and colleagues found differences in the duration of climax scream and the frequency range of the fundamental. While these acoustic features explained more variance than other features in our data, they were not significantly different among communities. We instead found statistically significant differences in the acoustic features related to noise (maximum and mean Wiener entropy coefficient) and in the maximum difference in peak frequency between successive time segments. The differences in noisiness in the acoustic features of the scream are less likely to be explained by vocal learning and more likely to be explained by other individual factors such as age, rank, and health. As Riede et al. (2007) suggest, producing loud and tonal climax screams is physiologically costly and hence, signals the physical condition of the caller. We therefore expect individuals that are in better physical condition (in terms of age and health) to exhibit lower noisiness in their climax screams (Fitch et al., 2002; Riede et al., 2004, 2007). Chimpanzees from Gombe in our sample were on average younger than chimpanzees from Kibale (median age at Kasekela: 20.5 (range: 16-33) years, Mitumba: 15 (range: 14-27) years, and Kanyawara: 25 (range: 15-55) years; Table 1). Hence, the observed differences between Gombe and Kibale chimpanzees could be attributed to age differences, but more detailed analyses are needed to confirm this.

In contrast to Crockford et al. (2004), we did not observe any significant differences among adjacent communities, and only observed a significant difference when we included a geographically distant community in the comparison. As Crockford and colleagues argued, a greater difference in neighboring communities compared to a geographically distant community would imply a greater role of vocal learning compared to other factors. This is because a greater difference in neighboring communities suggests that individuals are learning and matching calls to their community members to have more distinct vocalizations compared to their hostile neighbors. Any differences observed among geographically distant communities provides a baseline for the expected variation among communities that may be an outcome of factors other than vocal learning, such as habitat acoustics, sound environment, body size, and stochastic variation among individuals (Mitani et al., 1999). Given the geographical differences between Gombe and Kibale, the differences we observed may be explained by aforementioned factors, or simply due to the genetic differences expected among geographically distant communities. Hence, we did not find evidence of vocal learning in chimpanzee pant-hoots.

In contrast to communities and contexts, we found substantial differences among individuals. Individuals differed in structural features and in climax scream features, but not in build-up element features. When all features were taken together, we observed the strongest differences in the climax scream features. The temporal properties that revealed greatest individual distinctiveness were duration of the climax phase, duration from build-up to letdown, and number of climax elements and screams. The spectral acoustic features showing the greatest individual differences were acoustic features related to the fundamental frequency F0.

Specifically, the start, minimum, maximum, and mean F0, frequency range of F0, and minimum peak frequency were the features with the strongest individual level signal. Some of these acoustic features that correlated with individual differences were consistent with those identified by Crockford et al. (2004): maximum F0 and minimum peak frequency. Additionally, our results are consistent with results from previous studies by Mitani and colleagues (Mitani et al., 1999; Mitani & Brandt, 1994). Mitani & Brandt (1994) found that the PC1 that explained the most variance among individuals loaded most highly in acoustic features of the fundamental frequency F0 including, start, minimum, maximum, and mean F0. Similarly, Mitani et al. (1999) found significant individual differences in the minimum, maximum, and mean F0, and the frequency range of F0.

The consistencies and inconsistencies of our results with previous studies reveal several insights and raise new questions. Consistent with previous studies, our study confirms the individual distinctiveness of chimpanzee pant-hoots in both spectral and temporal properties. Our study also found some differences in the temporal properties of pant-hoots given in feeding and traveling contexts, confirming the possibility of some contextual encoding. In terms of community-specific differences, we could not confirm previous studies and only found differences in one category of acoustic features of geographically distant communities. Our findings support the views of Mitani et al. (1999), who proposed that factors other than vocal learning explain the observed differences among populations.

Our failure to find evidence for community-specific signatures could reflect features peculiar to Gombe chimpanzees, or alternatively, it may be the case that previous findings of differences among communities resulted from statistical artifacts. Several community-specific peculiarities can lead to differential selection pressures for community-specific vocalizations. For example, (i) a recent history of intergroup violence could lead to a greater selection pressure for community-specific vocalizations to facilitate identifying own community vs. neighbors. There is a history of lethal intergroup violence in Gombe (Wilson et al., 2004), Kibale (Watts et al., 2006), as well as in Taï chimpanzees studied by Crockford et al. (2004) (Boesch et al., 2008). However, Gombe chimpanzees have experienced a higher rate of inter-community killings (Boesch et al., 2008; Wilson et al., 2004), suggesting that the selection for community-specific vocalizations should be at least as strong as that for Taï chimpanzees, if not higher. (ii) Stability of hierarchy and strength of affiliative bonds in the community promote vocal convergence (Fedurek, Machanda, et al., 2013; Mitani & Brandt, 1994; Mitani & Gros-Louis, 1998) and thus could create positive selection pressure for community-specific vocalizations. In Gombe, within-community bonds are likely stronger in the Kasekela community, which has more maternal brothers (Bray & Gilby, 2020) compared to Mitumba, which has fewer brothers and higher within-community violence (Massaro et al., 2021). More data are needed to accurately test if social bonds affect vocal convergence across field sites. (iii) A larger community size may lead to a greater selection pressure for community-specific signatures as it becomes more difficult to keep track of individuals. All communities in this study and in Crockford et al. (2004) were relatively small, so this is less likely to explain the discrepancies in the results. Furthermore, while Crockford and colleagues attempted to control for confounding factors, their sample size of only three individuals per community increases the possibility that apparent differences could emerge by chance. Measurements common in studies of animal behavior tend to be inherently noisy and such noisy measurements are likely to exaggerate effect sizes and return false positives (Loken & Gelman, 2017). Our study included a greater number of individuals per community (5-7 individuals per community compared to 3 individuals per community in Crockford et al. (2004)) and hence should have detected any differences among communities that were similar in effect size to those reported by Crockford et al. (2004). Further, neither our study, nor Crockford et al. (2004) controlled for individual level factors such as age, body size, health condition, and rank that could influence the acoustic structure. In addition, no studies have been able to quantitatively control for other factors such as the influence of habitat differences and sound environments that Mitani et al. (1999) suggested could be important.

Our results reinforce the importance of replicating findings in animal behavior research. A key feature of scientific discovery is seeking results that are consistently reproducible (Burman et al., 2010; Johnson, 2002; Lamal, 1990; Popper, 1959). In recent decades, analysis of studies in several scientific disciplines, including fields as diverse as psychology and medicine, have found that most scientific findings fail to be reproduced by subsequent studies, leading to what has been called the replication crisis (Ioannidis, 2005; Wiggins & Chrisopherson, 2019). One factor contributing to this crisis is that studies replicating existing findings are rarely conducted, and are implicitly discouraged through reviewer bias against them (Neuliep & Crandall, 1993). Given that field studies in animal behavior typically have smaller sample sizes than studies in psychology or medicine, it is likely that the field of animal behavior is in even greater need of replication to test the validity of previous results with sufficient sample sizes (Johnson, 2002). Within animal behavior, the need for replication may be particularly acute for species such as chimpanzees, for which field conditions make it challenging to obtain sample sizes sufficient to be confident in results. Long-term data from multiple field sites have proven essential for providing sufficient sample sizes for a range of topics (*e.g*., culture: (Whiten et al., 1999); reproductive cessation: (Emery Thompson et al., 2007); lethal aggression: (Wilson et al., 2014)). Such collaboration across long-term studies will be essential for answering questions about vocal communication as well.

## Acknowledgements

We thank the Gombe Stream Research Centre field staff for carrying out the work needed to make research and conservation in and around Gombe possible, and the Jane Goodall Institute for supporting these projects. We thank Anthony Collins, Deus Mjungu, and Dismas Mwacha for additional support during the fieldwork. We thank the University of Minnesota for funding this research through the Talle Faculty Research Award to MLW. We thank Frans Plooij and Twise Victory BV for providing further funding to the Chimpanzee Vocal Communication Project at Gombe. We thank the Government of Tanzania, Tanzania National Parks, Tanzania Commission for Science and Technology, and Tanzania Wildlife Research Institute for permission to carry out the research. We thank Kurt Hammerschmidt, Roger Mundry, and Julia Fischer for providing advice and software packages to perform the acoustic and statistical analysis.

We thank the directors of the Kibale Chimpanzee Project (KCP), Richard Wrangham and Martin Muller, for allowing us to conduct this research on the Kanyawara chimpanzees. We thank the KCP field manager, Emily Otali, and KCP field assistants, Francis Mugurusi, Solomon Musana, James Kyomuhendo, Wilberforce Tweheyo, Sunday John, and Christopher Irumba, who were extremely helpful during the fieldwork. PF was funded by a BBSRC studentship, a Leakey Foundation General Grant, and an American Society of Primatologists General Small Grant. KS was funded by a BBSRC project grant.

