## Supplementary materials for "Chimpanzee pant-hoots encode information about individual but not group differences"

Figure S1 (a): Buildup to letdown duration at individual and community levels.

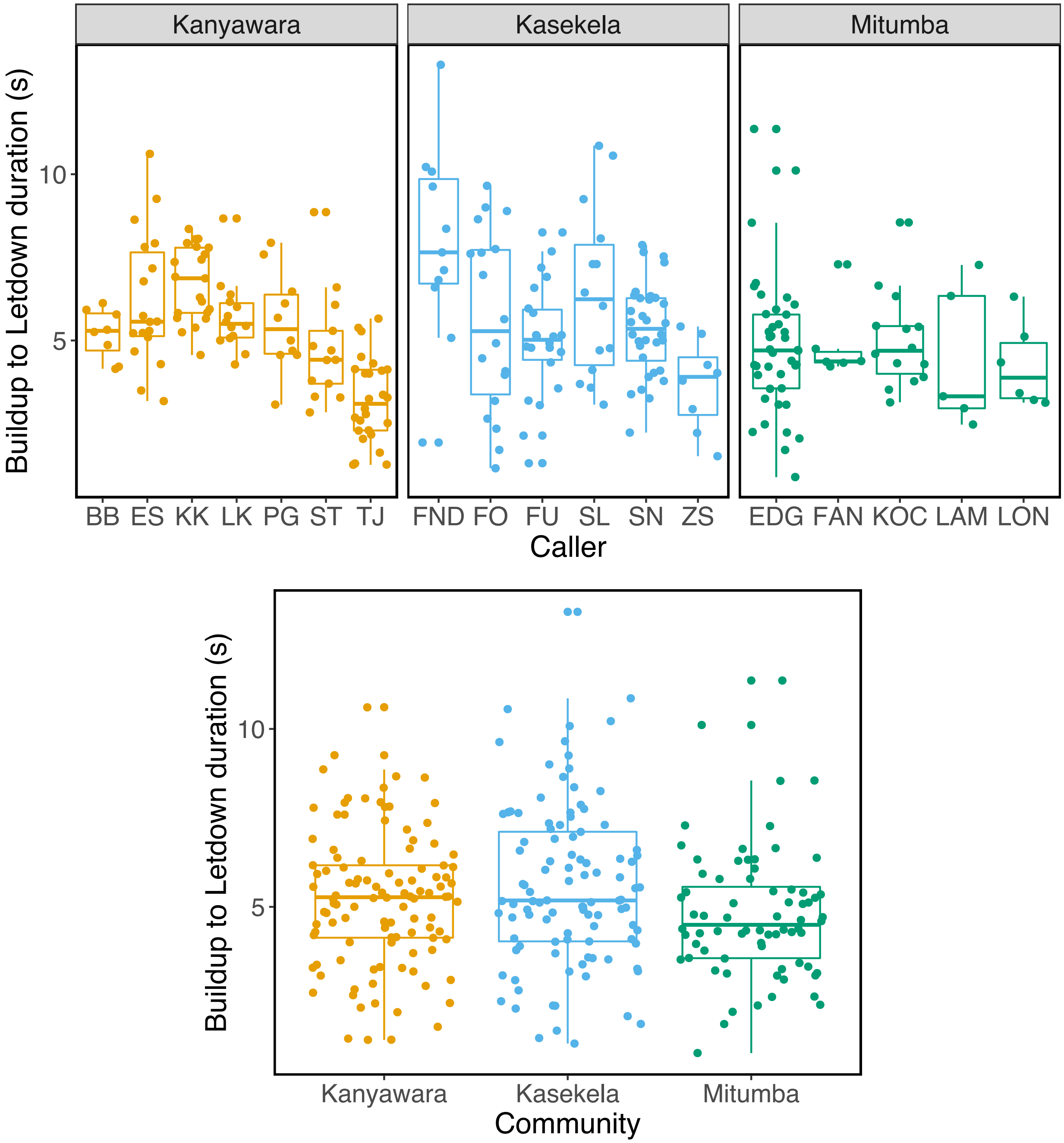

Figure S1 (b): Proportion of calls with build-up at individual and community levels.

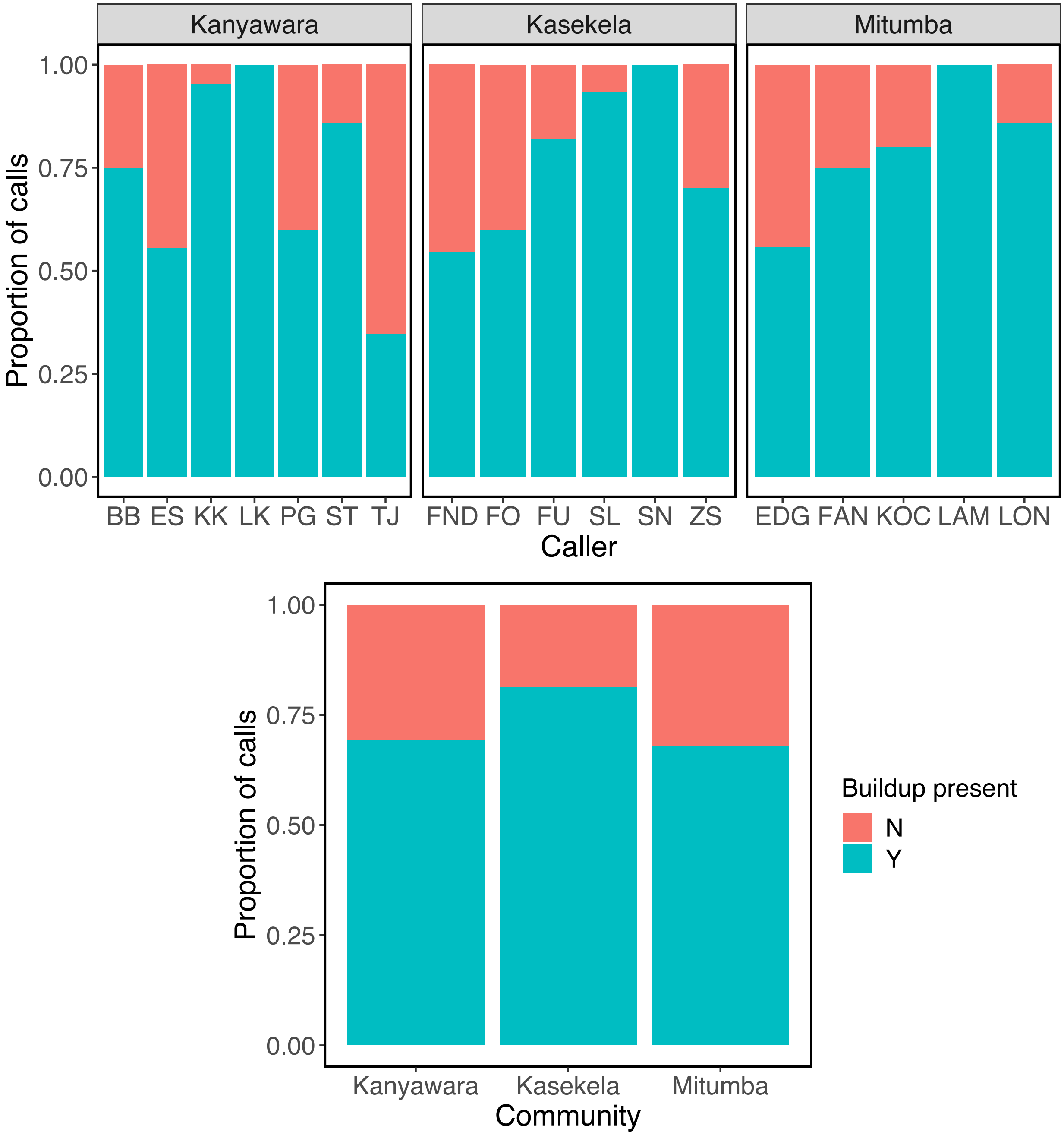

Figure S1 (c): Build-up duration at individual and community levels.

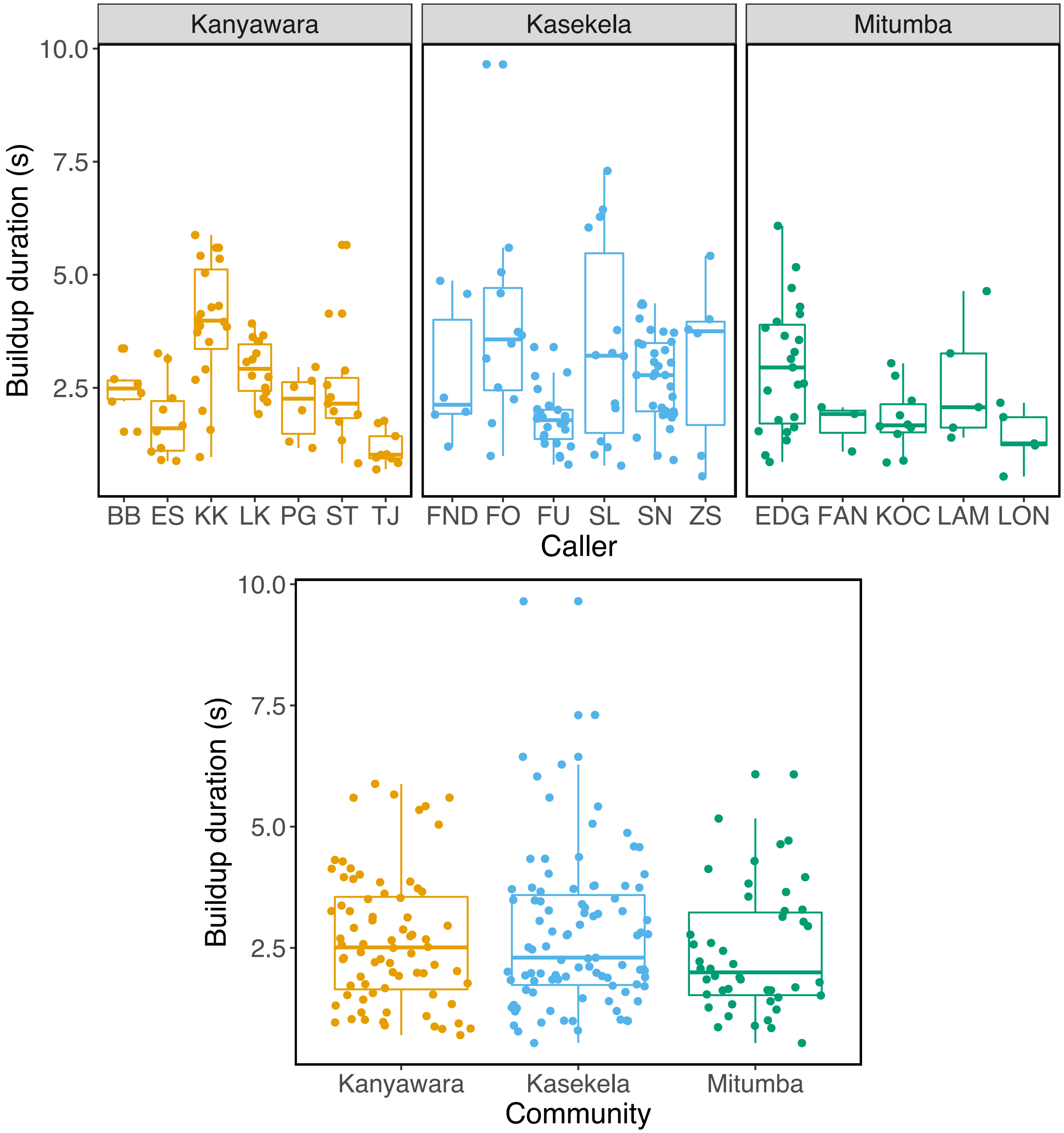

Figure S1 (d): Number of build-up exhalation components at individual and community levels.

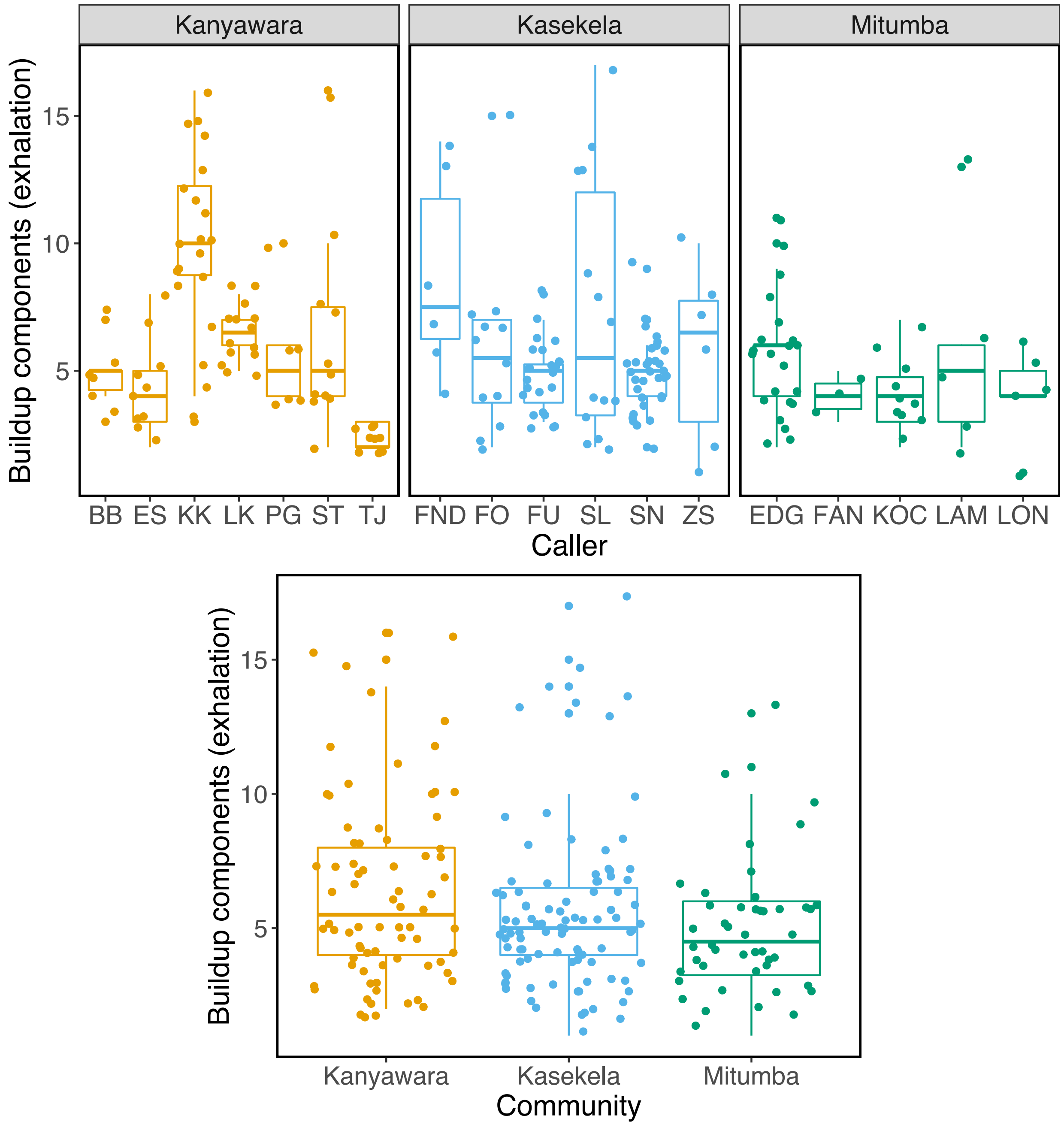

Figure S1 (e): Rate of build-up at individual and community levels.

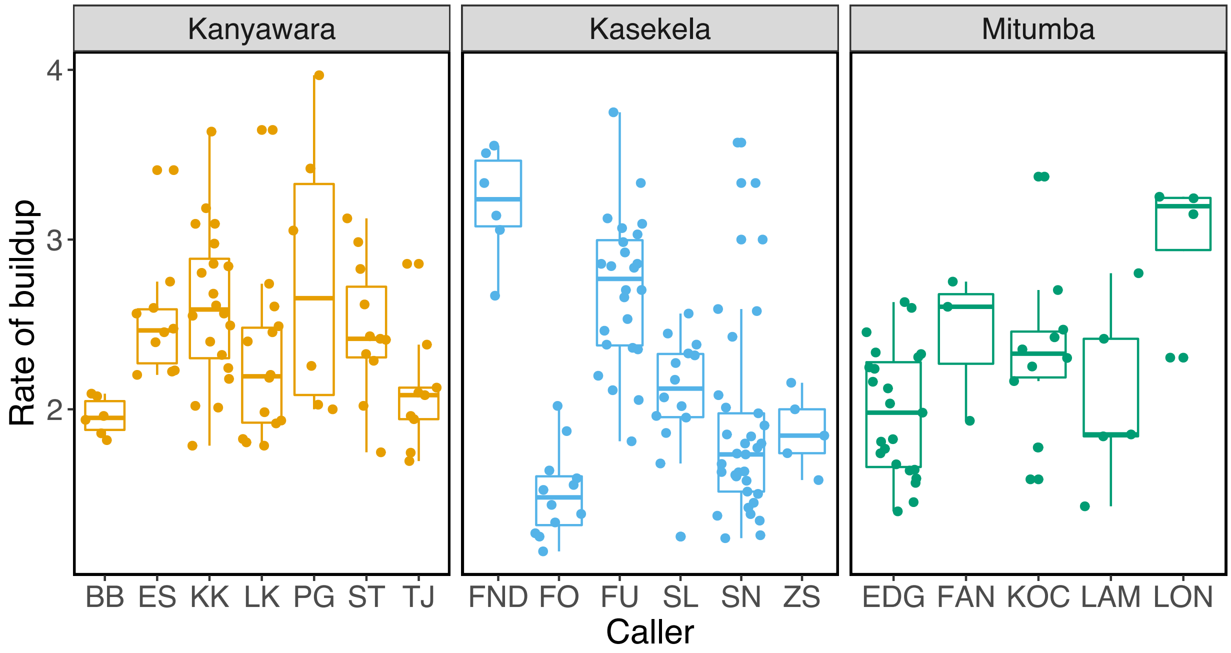


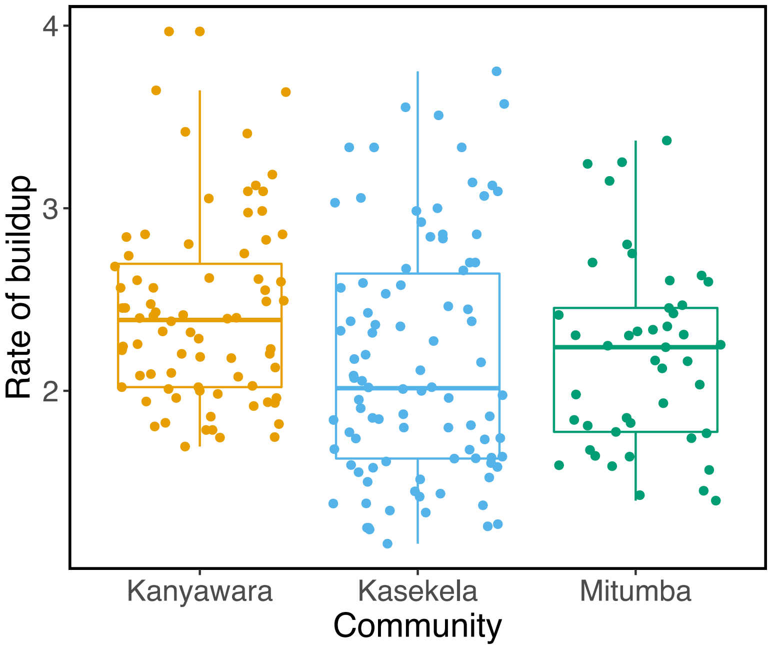

Figure S1 (f): Climax duration at individual and community levels.

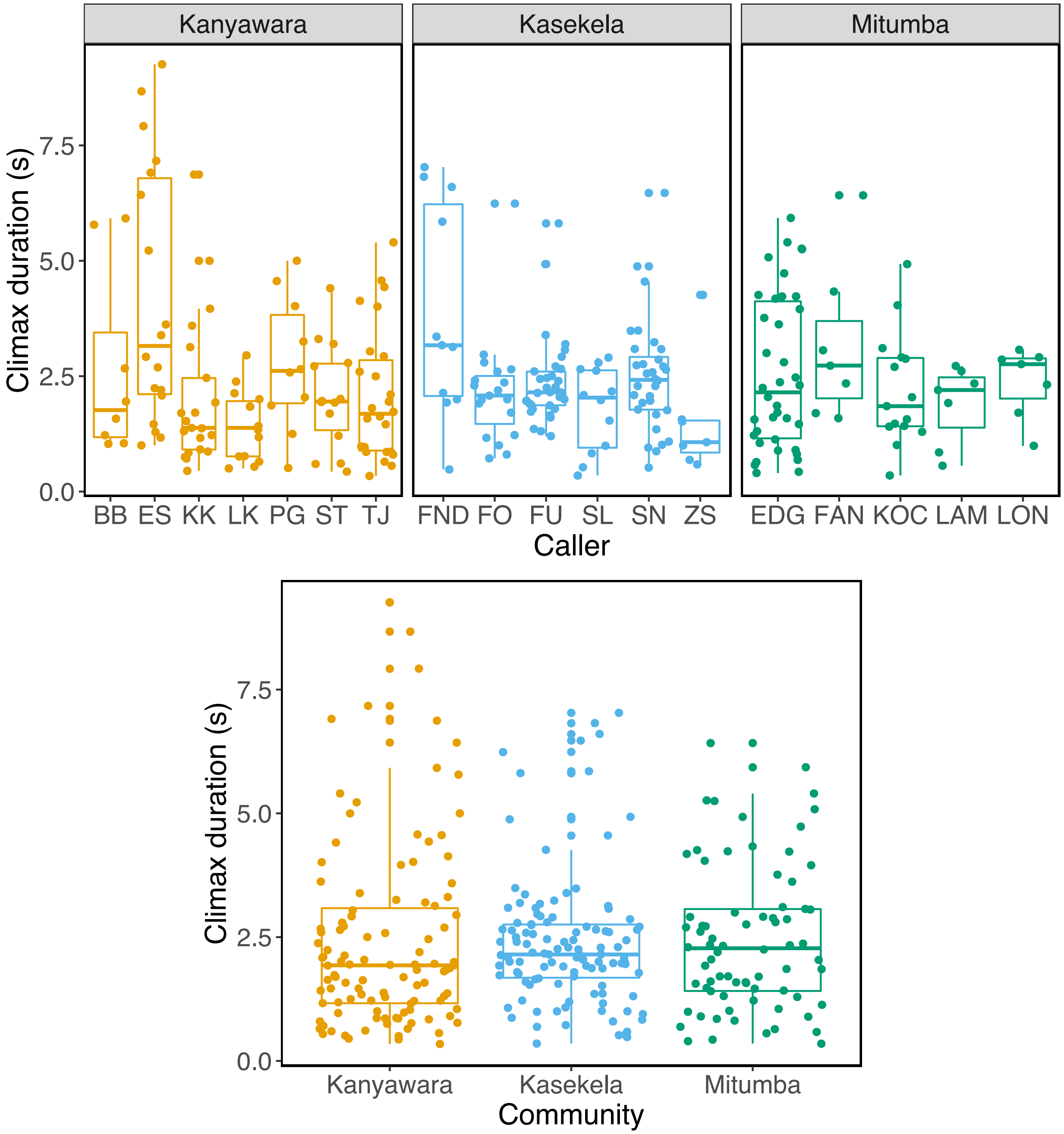

Figure S1 (g): Number of climax components at individual and community levels.

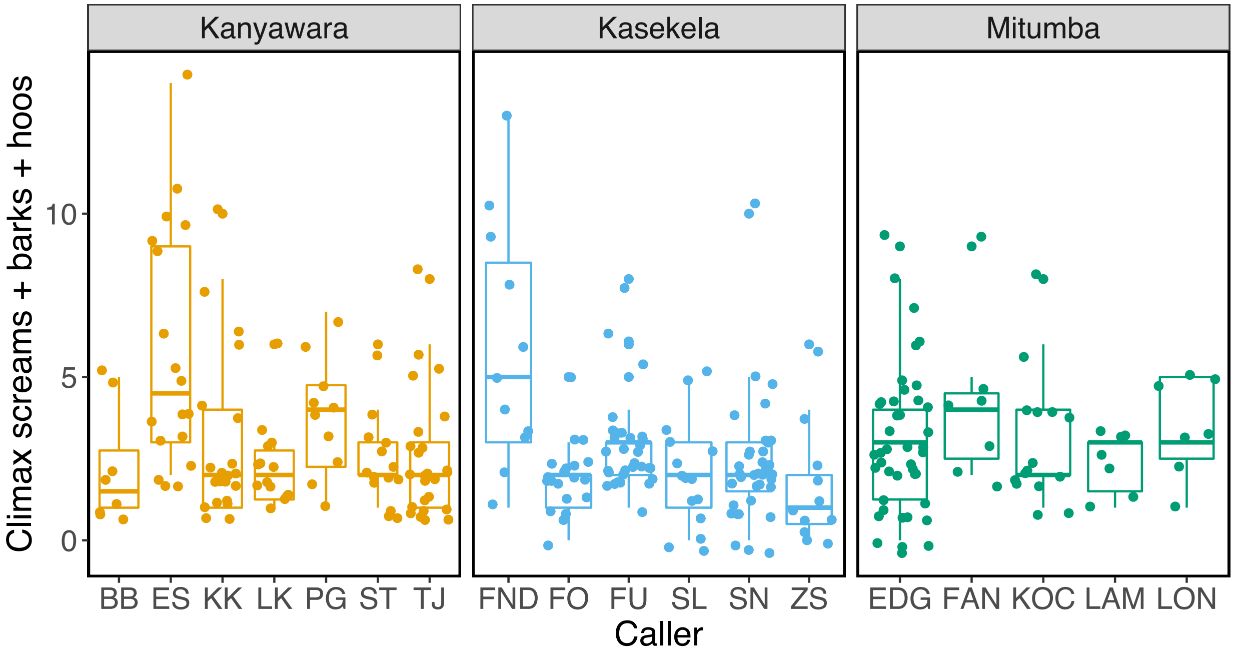


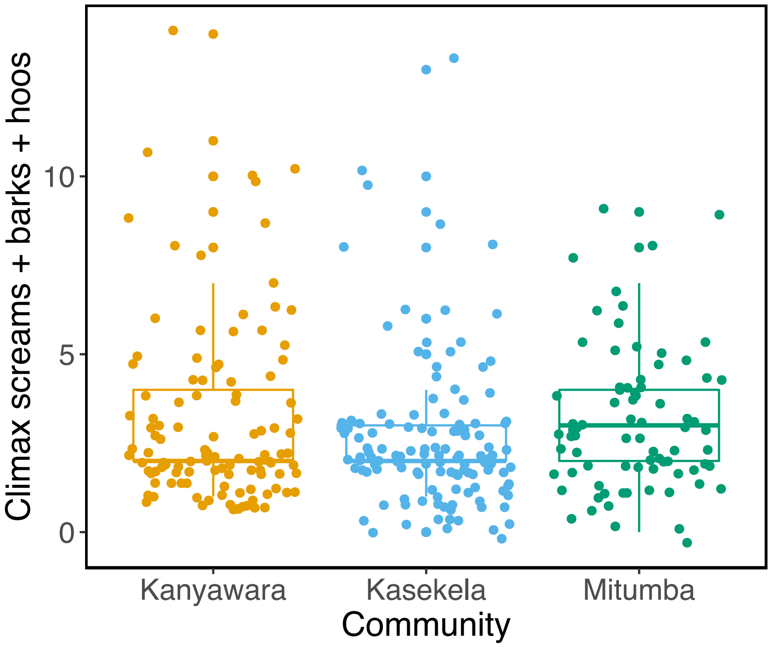

Figure S1 (h): Number of climax screams at individual and community levels.

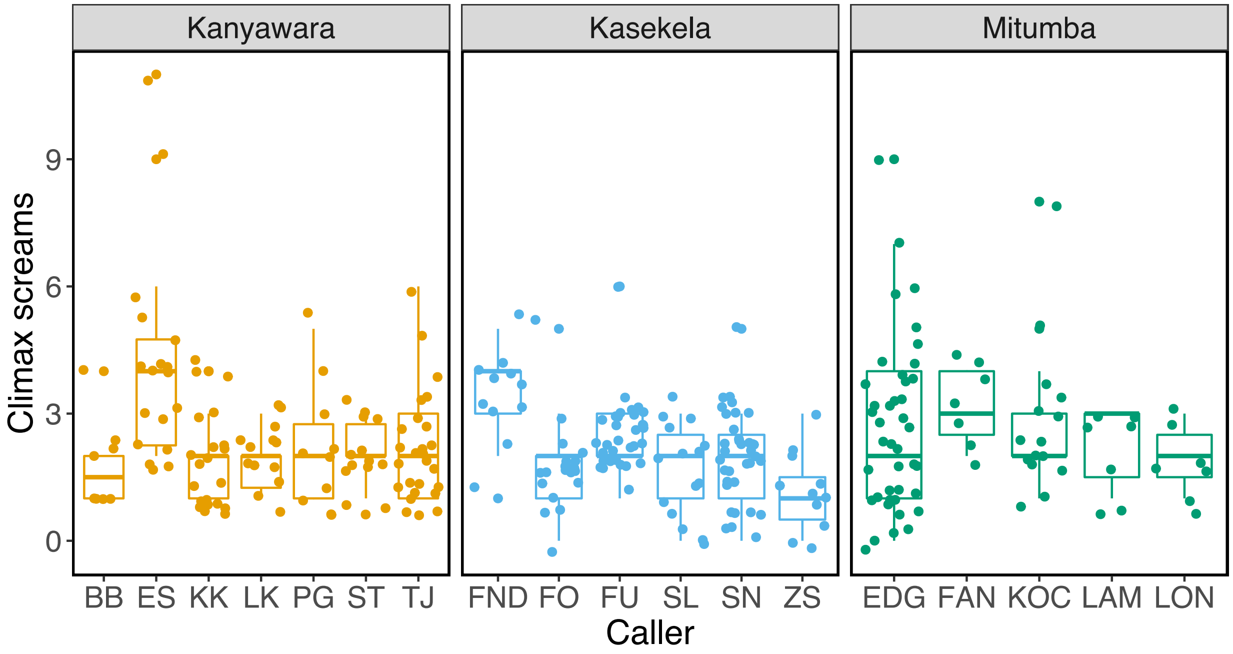


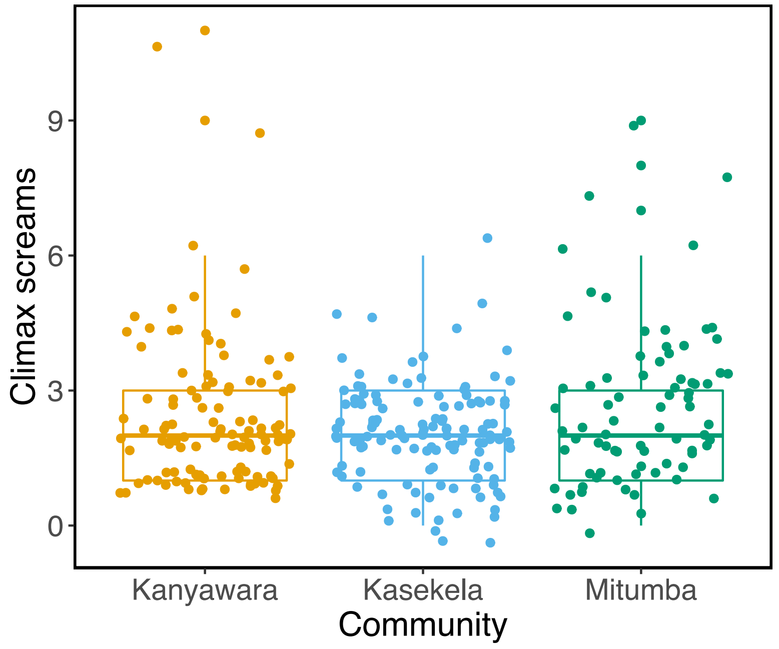

Figure S1 (i): Proportion of climax components that are screams at individual and community levels.

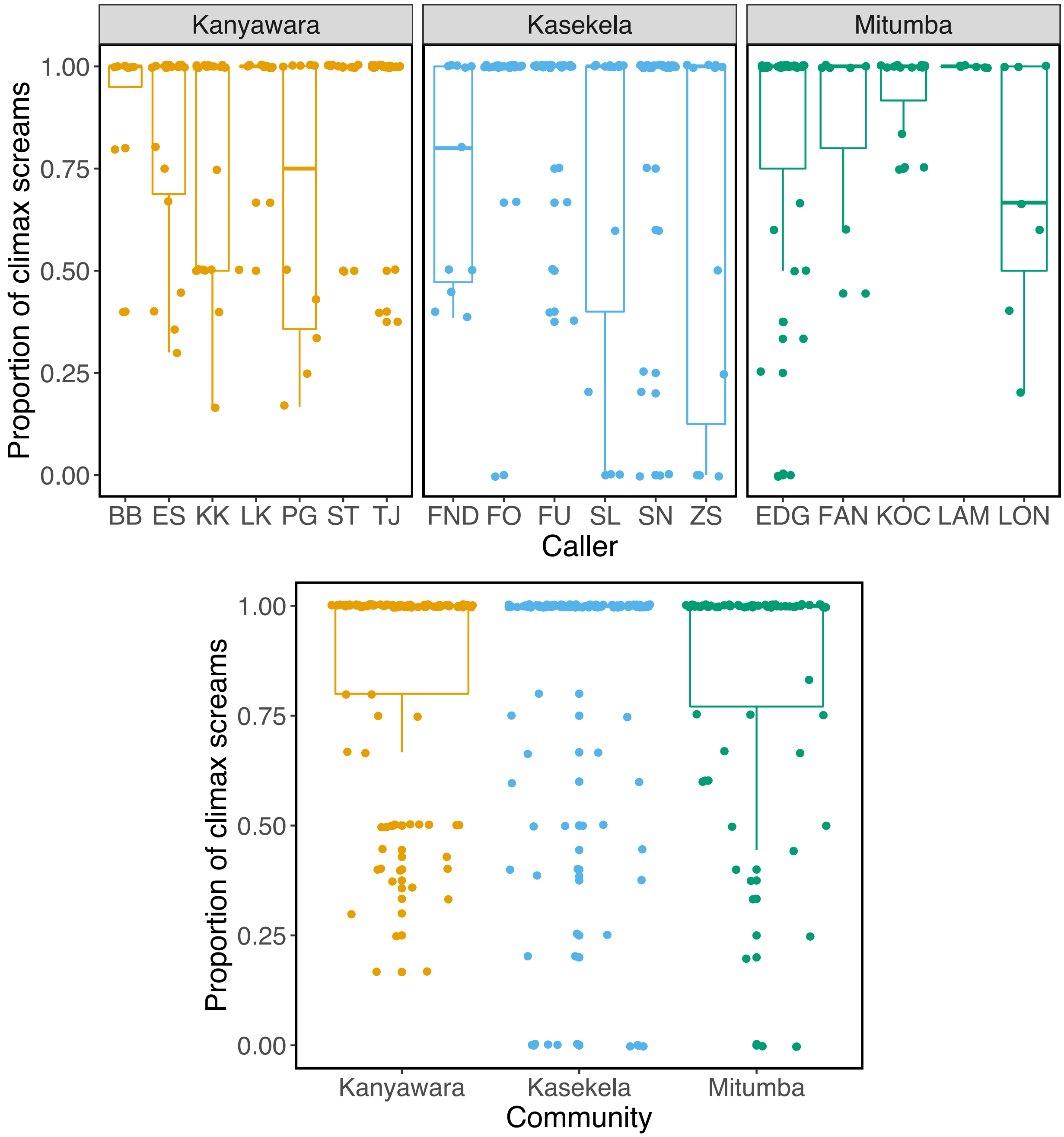

Figure S1 (j): Proportion of calls with letdown present at individual and community levels.

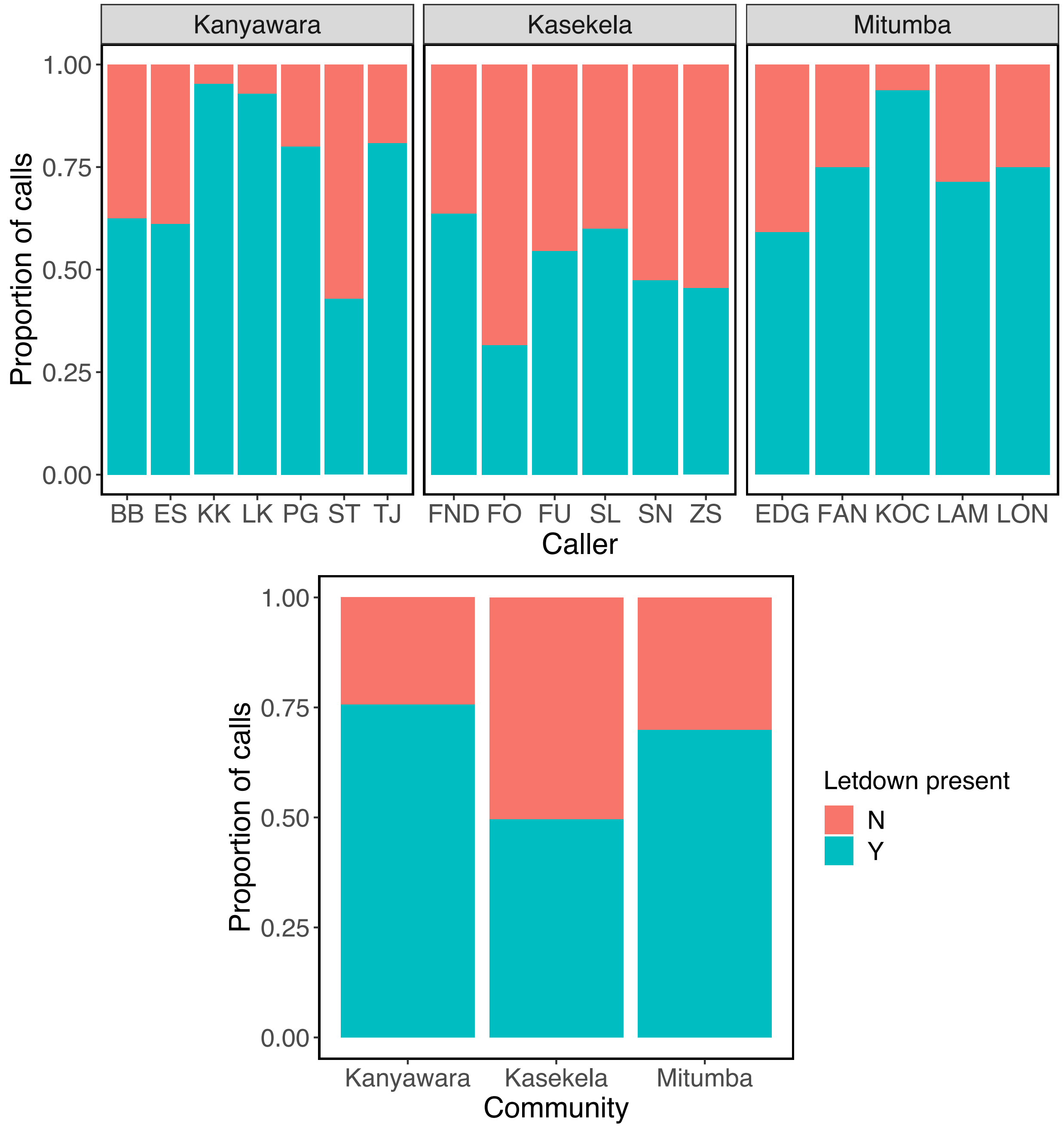

Figure S1 (k): Number of letdown components at individual and community levels.

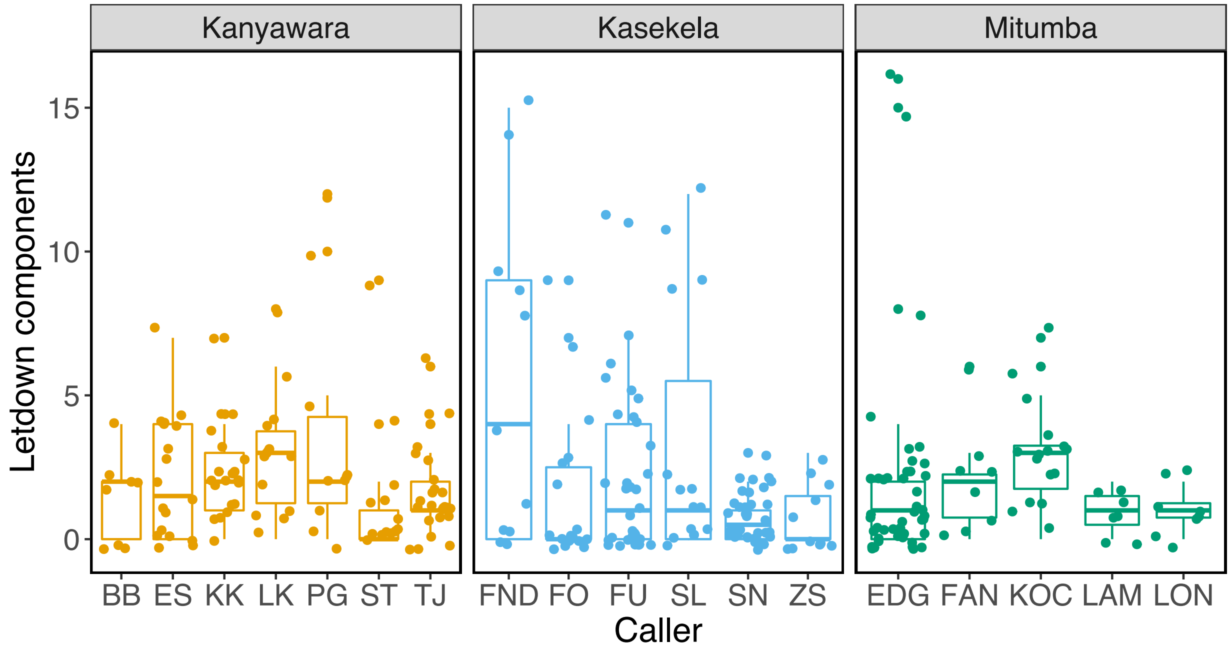


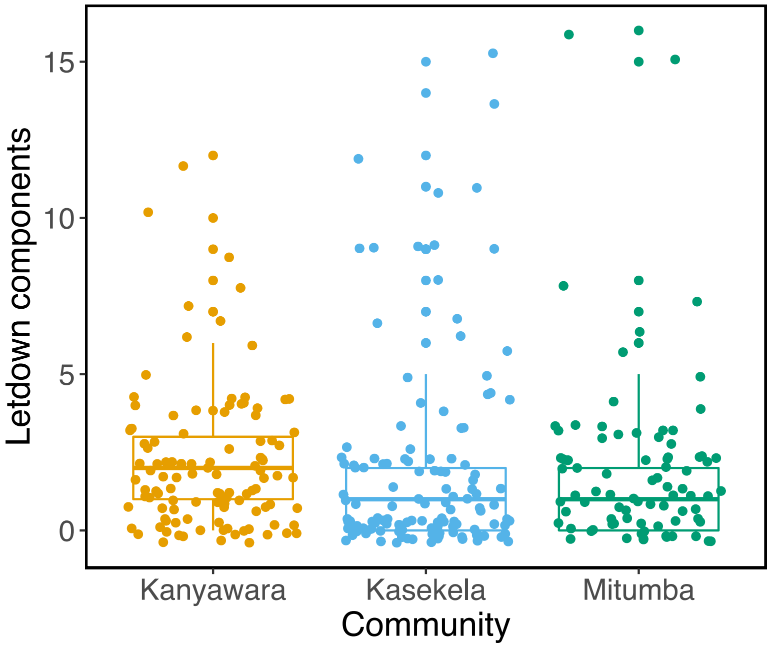

Figure S1 (l): Proportion of calls with drumming present at individual and community levels.

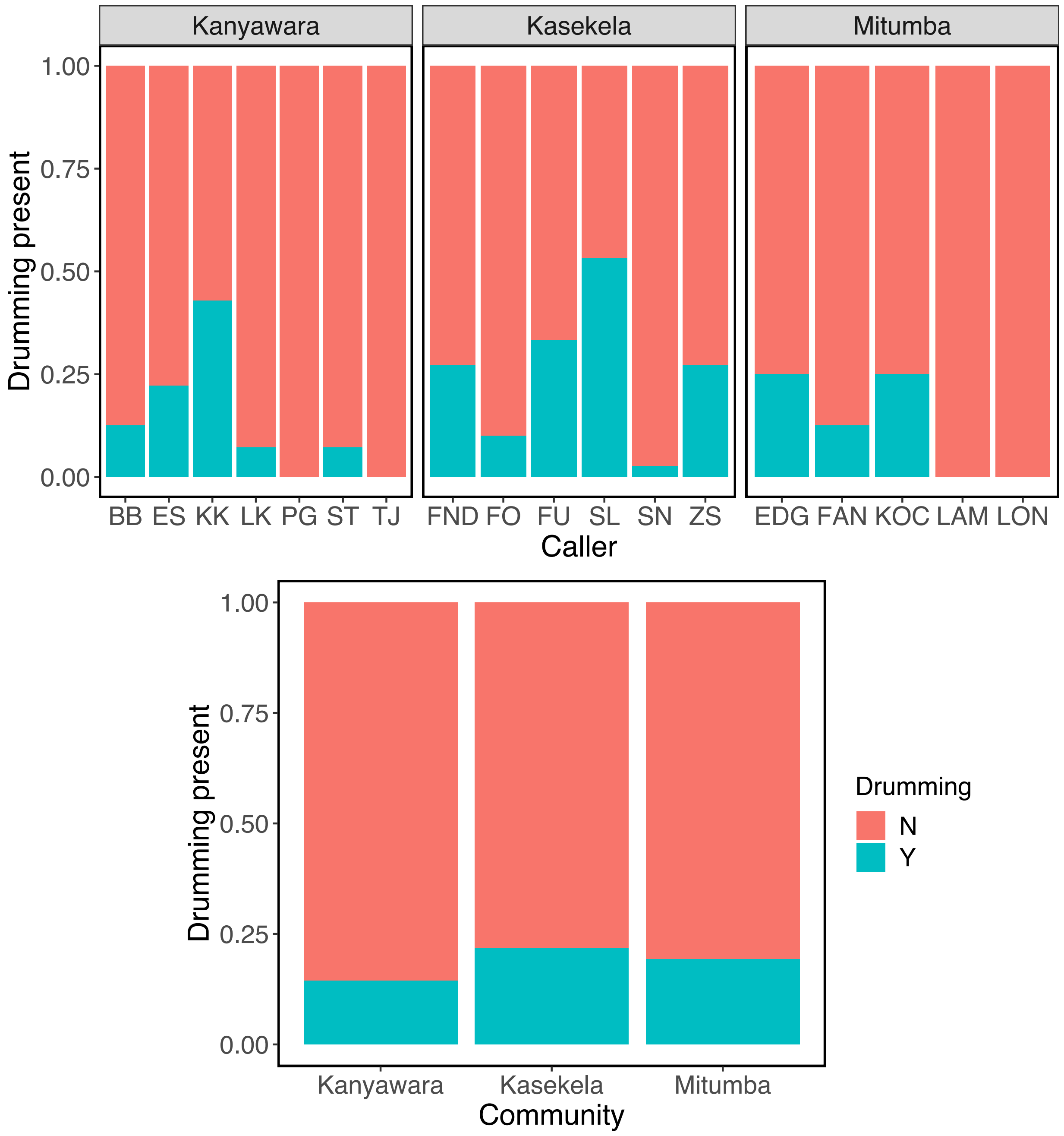

Figure S1 (m): Number of drum beats at individual and community levels.

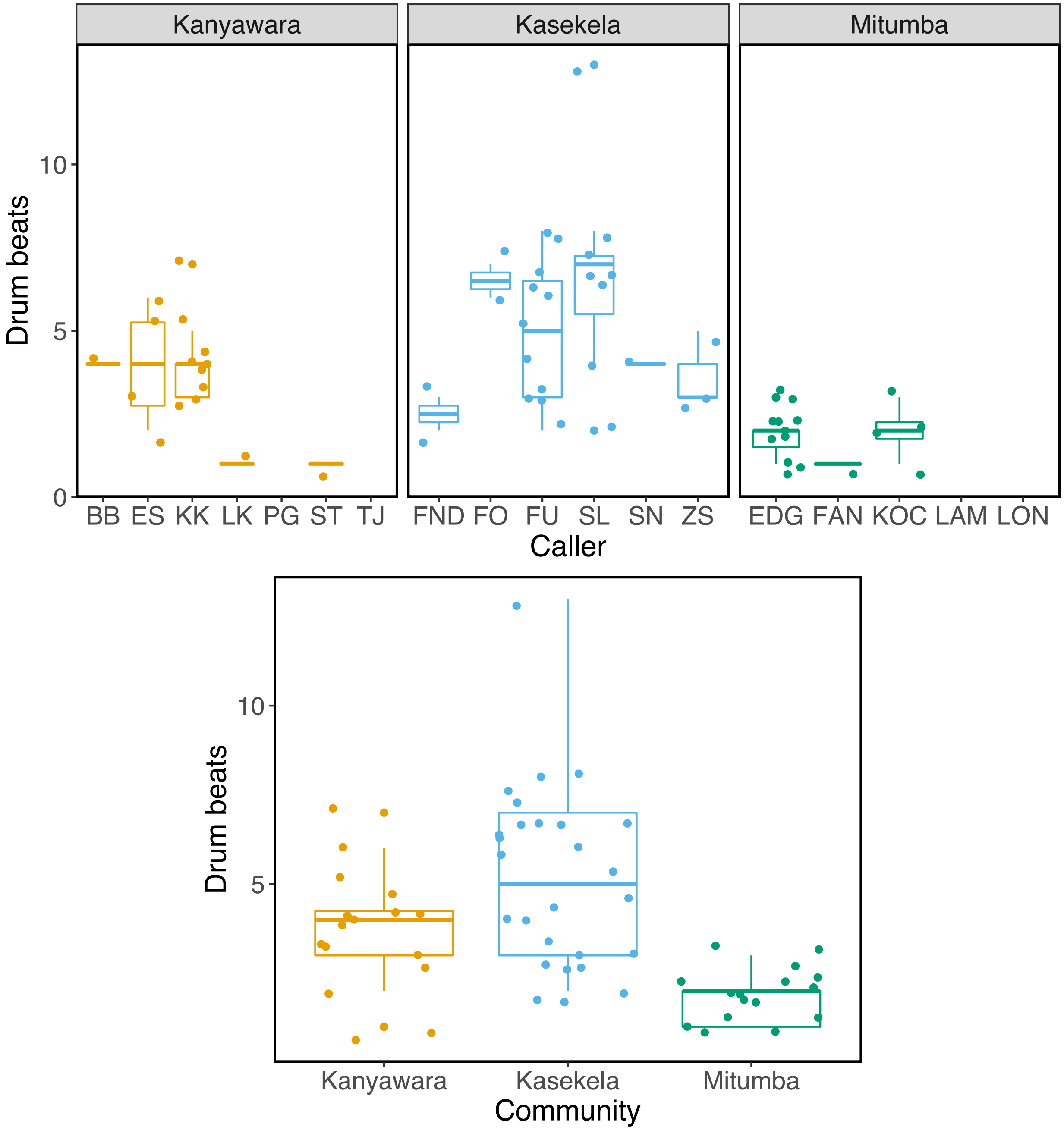

Figure S2 (a): Differences in the number of letdown components between contexts at individual and community levels.

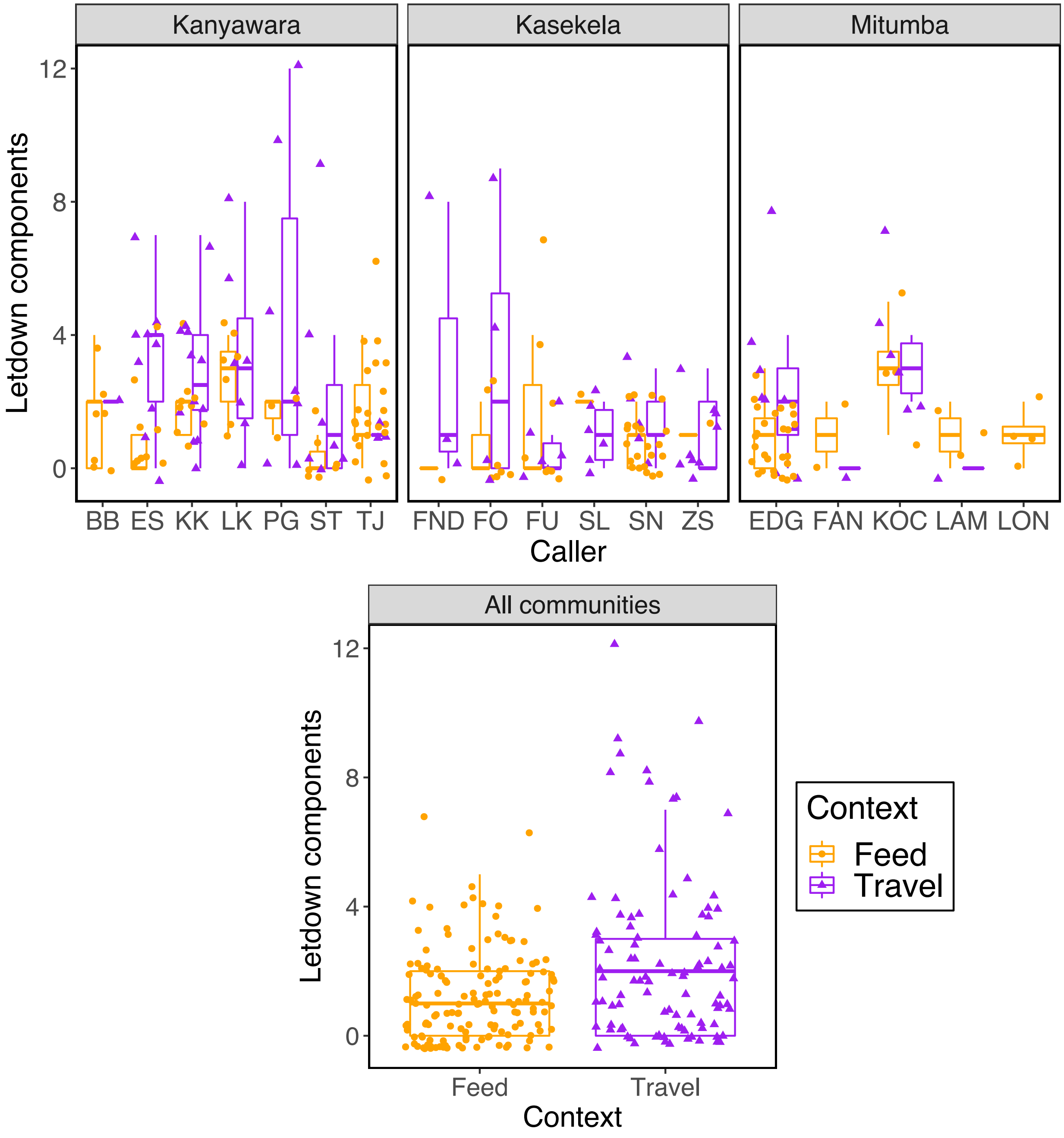

Figure S2 (b): Differences in the number of build-up components between contexts at individual and community levels.

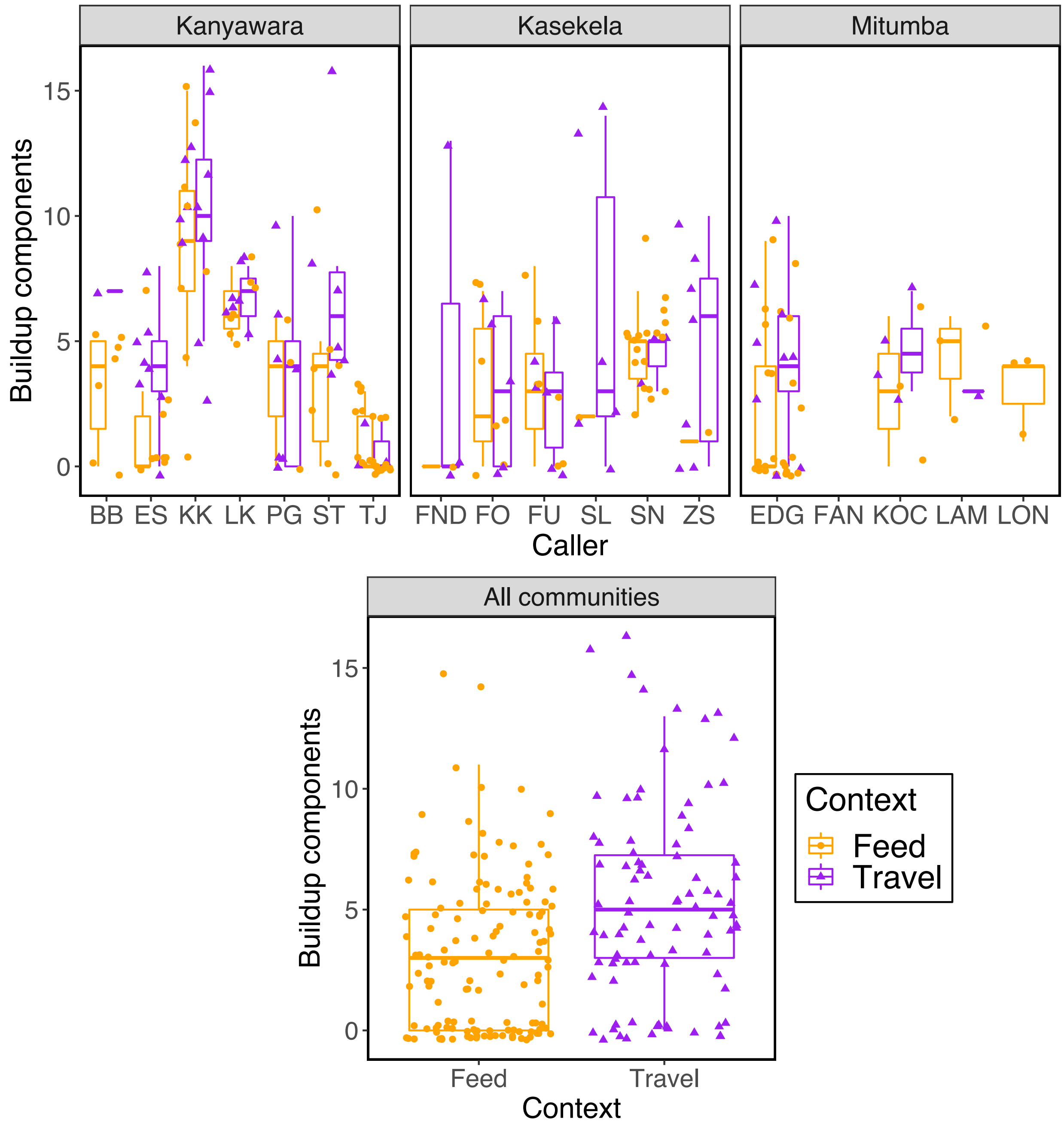

Figure S2 (c): Differences in the number of climax components between contexts at individual and community levels.

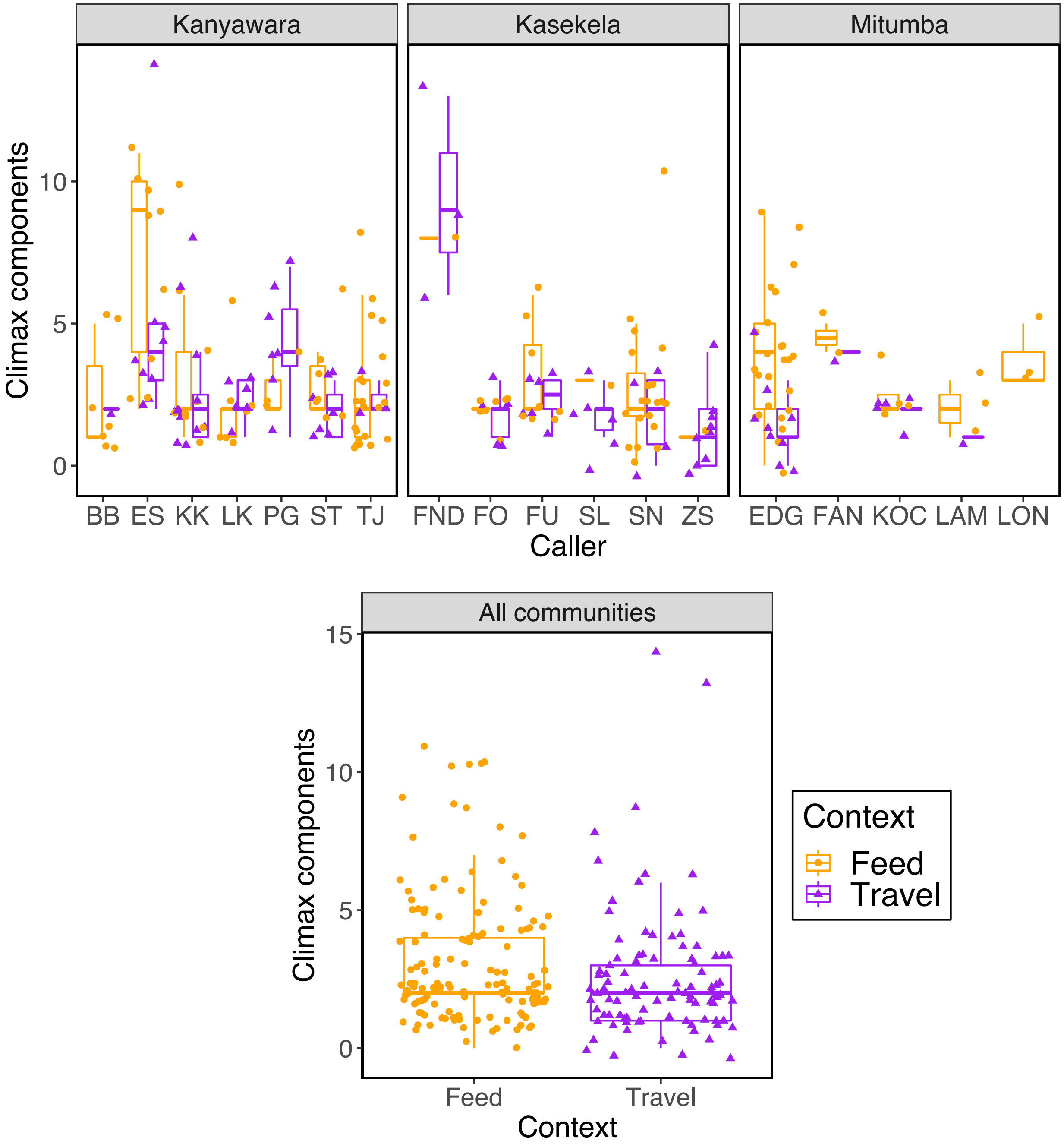
